# ClpP/ClpX deficiency impairs mitochondrial functions and mTORC1 signaling during spermatogenesis and meiosis

**DOI:** 10.1101/2022.12.16.520702

**Authors:** Chenxi Guo, Yuan Xiao, Jingkai Gu, Zhe Hu, Jiahuan Zheng, Renwu Hua, Zhuo Hai, Jiaping Su, Jian V. Zhang, William S.B. Yeung, Tianren Wang

**Affiliations:** Shenzhen Key Laboratory of Fertility Regulation, Reproductive Medicine Center, The University of Hong Kong-Shenzhen Hospital; Shenzhen, 518053, China; Department of Obstetrics and Gynaecology, Li Ka Shing Faculty of Medicine, The University of Hong Kong; Hong Kong SAR, 999077, China; Center for Energy Metabolism and Reproduction, Shenzhen Institute of Advanced Technology, Chinese Academy of Sciences; Shenzhen, 518055, China; Shenzhen Key Laboratory of Metabolic Health; Shenzhen, 518055, China

## Abstract

Caseinolytic protease proteolytic subunit (ClpP) and caseinolytic protease X (ClpX) are mitochondrial matrix peptidases that activate mitochondrial unfolded protein response to maintain protein homeostasis in the mitochondria. However, the role of ClpP and ClpX in spermatogenesis remains largely unknown. In this study, we demonstrated the importance of ClpP/ClpX for meiosis and spermatogenesis with two conditional knockout (cKO) mouse models. We found that ClpP/ClpX deficiency reduced mitochondrial functions and quantity in spermatocytes, affected energy supply during meiosis and attenuated zygotene-pachytene transformation of the male germ cells. The dysregulated spermatocytes finally underwent apoptosis resulting in decreased testicular size and vacuolar structures within the seminiferous tubules. We found mTORC1 pathway was over-activated after deletion of ClpP/ClpX in spermatocytes. Long-term inhibition of the mTORC1 signaling via rapamycin treatment *in vivo* partially rescue spermatogenesis. The data reveal the novel roles of ClpP and ClpX in regulating meiosis and spermatogenesis.

**One-Sentence Summary:** ClpP/ClpX is required for maintaining mitochondrial functions in spermatocytes during meiosis and spermatogenesis.

## INTRODUCTION

Mitochondria are widely known to play a central role in cellular energy production and supply (*1*). The chemical energy produced by oxidative phosphorylation in the mitochondria is stored in adenosine triphosphate (ATP) (*2*). The energy generated by mitochondria are crucial for various cellular activities, including gene transcription, protein synthesis, cell differentiation and growth, mitosis and meiosis (*3-7*). Mitochondria also provide a platform for metabolic pathways, including tricarboxylic acid cycle, oxidation, and lipid synthesis (*8*). The functional versatility and adaptability of mitochondria are based on numerous proteins, protein complexes, as well as stress-response pathways (*9*). A mature mammalian spermatozoon contains around 80 mitochondria in the mid-piece (*10*). In addition to energy production, mitochondria in spermatozoa are implicated in redox equilibrium and calcium regulation essential for capacitation, sperm motility, acrosome reaction and gamete fusion (*11*). Available evidences suggest that mitochondria also play diverse roles in germ cell differentiation and spermatogenesis (*12, 13*). During spermatogenesis, the mitochondria change their morphology, size and location (*14*). The spermatogonia contain evenly distributed ovoid shaped mitochondria with lamellar cristae and an electron translucent matrix. The zygotene and early pachytene spermatocytes contain increasing number of elongated mitochondria, signifying an active functional state, in close proximity to the nuclear membrane (*14, 15*). The late spermatocytes and early spermatids contain mitochondria with enlarged electron-dense matrix and vesicular cristae, evenly distributed throughout the cytoplasm (*16*). However, the exact roles of mitochondria in spermatogenesis remain largely unknown. ClpP and ClpX are mitochondria-specific quality control proteases. They maintain proteostasis via degrading misfolded or damaged proteins in mitochondrial matrix (*17*). Maintaining protein homeostasis is a challenging work for the mitochondria. Disruption to proteostasis is usually caused by local reactive oxygen species (ROS) produced during normal and attenuated oxidative phosphorylation (Curtis et al., 2012). The ClpP and ClpX complex (ClpXP complex) is a AAA+ protease that uses the energy of ATP binding and hydrolysis to degrade unfolded or misfolded proteins (*18*). The complex consists of a hexamer of ClpX and a tetradecamer of ClpP (*19*). ClpX recognizes the protein substrates via binding with their unstructured peptide sequences, termed as degradation tags or recognition signals (*20*). ClpP acts as a proteolytic component of ClpX or other AAA+ proteases; ClpP binds to its partner ATPase and cleaves any polypeptide that is translocated to the proteolytic chamber of ClpP. The small fragments of the cleaved polypeptides can then exit the chamber and be further degraded by exopeptidases to free the amino acids of the fragments (*19, 21*).

Recent studies investigated the role of ClpP and ClpX in germ cell development and reproductive diseases (*22-24*). A case study reported a homozygous missense mutation of ClpP in an azoospermic man (*25*). ClpP mutations also cause the autosomal recessive Perrault syndrome (PRLTS), which is characterized by primary ovary insufficiency in females and early-onset permanent hearing loss in males and female (*26, 27*). These observations associate ClpP deficiency with blockage of germ cell differentiation and development, though the underlying mechanism is unknown. The research on ClpX is limited. ClpX can exist as an ATP-dependent molecular chaperone alone, rather than as a ClpXP complex in some species, like yeast, indicating that the ClpXP complex is dispensable in some organisms (*28*). The role of ClpX in germ cell differentiation and development awaits further investigation.

Here, we utilized the Cre-LoxP system and generated conditional knock-out (cKO) mouse lines with specific knock-out of ClpP or ClpX in the postnatal germ cells. Examination of the histology and structure of the testis in these mice showed that the deletion attenuated zygotene-pachytene transformation of the spermatocytes. We performed high-throughput sequencing on highly purified spermatocytes and found that the mTOR signaling was significantly altered after genetic deletion of the *Clpp* or *Clpx* gene. Interestingly, *in vivo* inhibition of mTORC1 can partially rescue the phenotypes. Overall, our data provide important insights into mitochondrial quality control in meiosis and spermatogenesis, and reveal a novel role of ClpP/ClpX in spermatogenesis via regulating the mTORC1 signaling.

## RESULTS

### ClpP or ClpX deficiency decreases testicular size and disrupts spermatogenesis

The ClpP/ClpX-floxed and Stra8-Cre (S8-Cre) cKO mice were genotyped (Fig. 1B). The higher band in the top panel represents a successful loxP insertion in the ClpP/ClpX genome, while the lower band represents the wild-type (WT) ClpP/ClpX band (Fig. 1Bi and Bii). Here, we utilized *ClpP*^*fl/fl*^ or *ClpX*^*fl/fl*^ animals as the WT controls. The *ClpP*^*fl/fl*^;*S8-Cre* or *ClpX*^*fl/fl*^;*S8-Cre* mice were termed ClpP cKO or ClpX cKO mice, respectively hereafter. We sacrificed animals from the same littermate for comparison in majority of experiments. The levels of ClpP and ClpX protein in the purified spermatocytes were determined. Western blotting analysis (Fig. 1Ci) showed that the ClpP cKO spermatocytes had undetectable level of ClpP and significantly higher expression of ClpX relative to the control group (Fig. 1Cii). Similarly, the spermatocytes of the ClpX cKO mice had almost no ClpX expression, but the expression of ClpP was significantly higher than the control (Fig. 1Ciii). Thus, the data not only demonstrated deletion of the ClpP or ClpX in the respective cKO mice, but also compensation of the ClpP and the ClpX protein after deletion of ClpX and ClpP, respectively. We next checked the testicular size of two cKO mouse lines. The sizes of the testis from both cKO mice were dramatically decrease at PD 56 when compared with the controls (Fig. 1Di and Dii). No difference was detected in body weight between the control and the ClpP/ClpX cKO animals (Fig. 1Ei), while the testis/body weight ratio showed significant decrease in both cKO mice relative to the control mice (Fig. 1Eii).

**Fig. 1.**
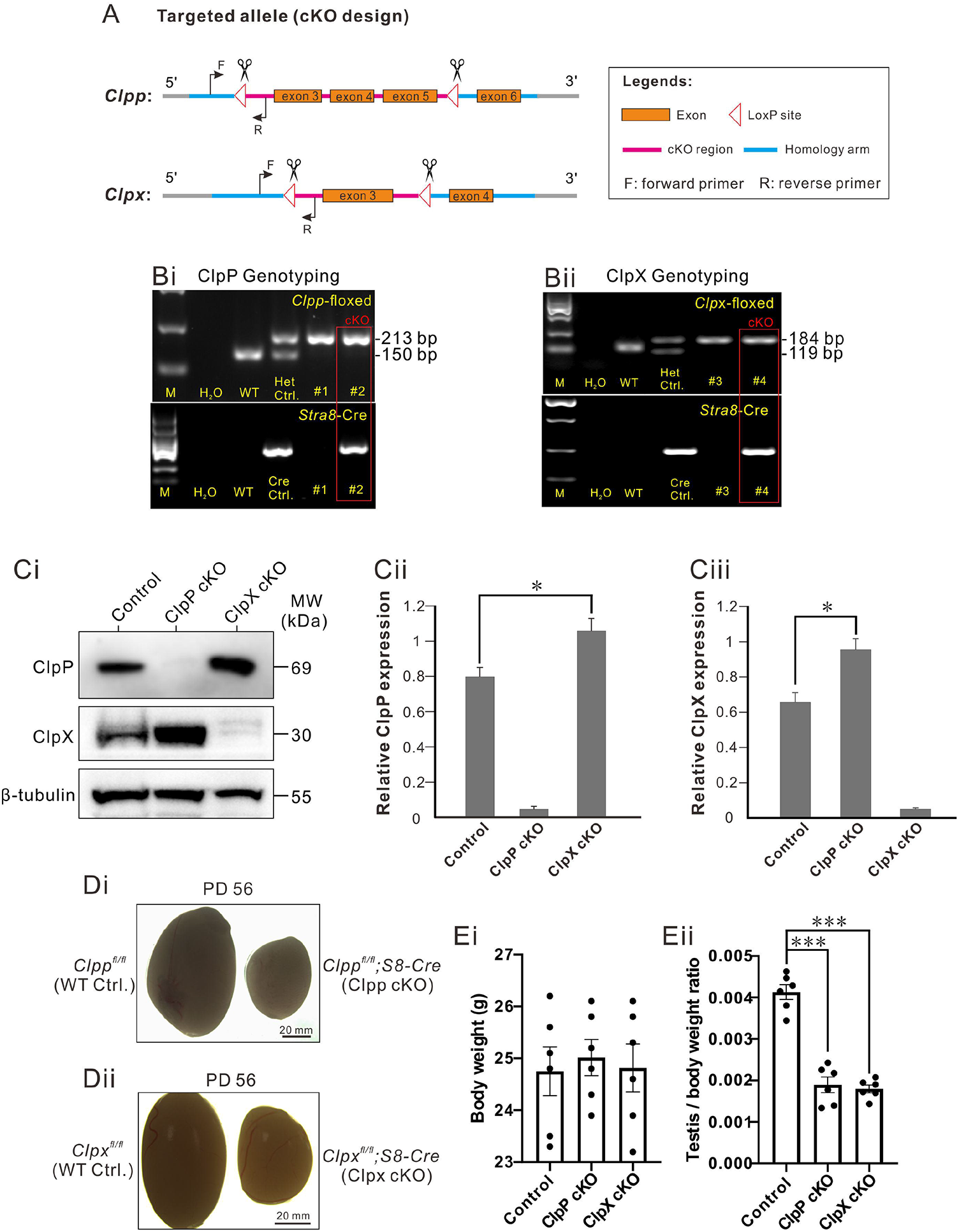
ClpP/ClpX conditional knock-out (cKO) mice demonstrate decreased testis size. (A) Design of the Cre-LoxP cKO system in the *ClpP/ClpX* allele, such that exon 3-5 was flanked by two LoxP regions in the *ClpP* allele, and exon 3 was flanked by two LoxP regions in the *ClpX* allele. When combined with Cre protein, exon 3-5 is deleted to complete *ClpP* gene knock-out; When combined with Cre protein, exon 3 is deleted to complete *Clpx* gene knock-out. The forward and reverse primers were designed to check the insertion of the LoxP site in the *ClpP/ClpX* allele. (Bi and Bii) Genotyping of the *ClpP*^*fl/fl*^*/ClpX*^*fl/fl*^; *Stra8-Cre* cKO mice. Tail DNA from wild-type (WT) mice was used as a negative control for PCR data analysis, whereas that from the *ClpP*^*fl/+*^*/ClpX*^*fl/+*^ heterozygous mouse was used as a positive control as there was a high band (213 base pairs, indicating successful insertion of LoxP in one of the *ClpP* alleles; 184 base pairs, indicating successful insertion of LoxP in one of the *ClpX* alleles) and a low band (150 base pairs, indicating the *ClpP* WT allele, without LoxP insertion; 119 base pairs, indicating the *ClpX* WT allele, without LoxP insertion). Tail DNA from the *Stra8-Cre* mouse was used as a positive control to amplify a Cre band for PCR analysis. (Ci) Western blot analysis to show the level of ClpP/ClpX expression in the spermatocytes of control and ClpP/ClpX cKO mice. β-tubulin was used as a loading control. The expression of (Cii) ClpP and (Ciii) ClpX were quantified in three groups. (Di,Dii) Representative examples of the testes isolated from *ClpP*^*fl/fl*^*/ClpX*^*fl/fl*^ mice and *ClpP*^*fl/fl*^*/ClpX*^*fl/fl*^; *Stra8-Cre* cKO mice, which were collected on postnatal day (PD) 56. The scale bar is 20 mm. (Ei,Eii) Bar charts to show the mean ± SEM (Ei) body weight and (Eii) testis weight/body weight ratio in the control and ClpP/ClpX cKO mice groups (n=6 for each group) at PD 56. *: p < 0.05, ***: p < 0.001.

Histological analysis of the testis and epididymis tissue from mice at PD 56 demonstrated strong negative effect on spermatogenesis in seminiferous tubules of the mutant animals (Fig. 2A-B). Figure 2Ai-Aiv shows images of overall patterns of testicle structure from the control and the mutant mice at low magnification. Severe germ cell loss was observed in majority of the seminiferous tubules in the ClpP cKO testis (Fig. 2Aii). The cells of the seminiferous tubules were not well organized with loose patches of germ cells and a lot of empty space in the tubules (Fig. 2Avi). In contrast to the tubules of the controls (*Clpp*^*fl/fl*^ or *Clpx*^*fl/fl*^) with germ cells at all stages of spermatogenesis (Fig. 2Av and Avii), the tubules of the ClpP cKO contained only spermatogonium (SG) and spermatocytes (SC), and almost without round spermatids (ST) and elongated spermatozoa (SZ). The presence of empty areas (red asterisks) in the central and peripheral regions of the seminiferous tubules indicated germ cell loss after ClpP germ cells specific deletion (Fig. 2Avi).

**Fig. 2.**
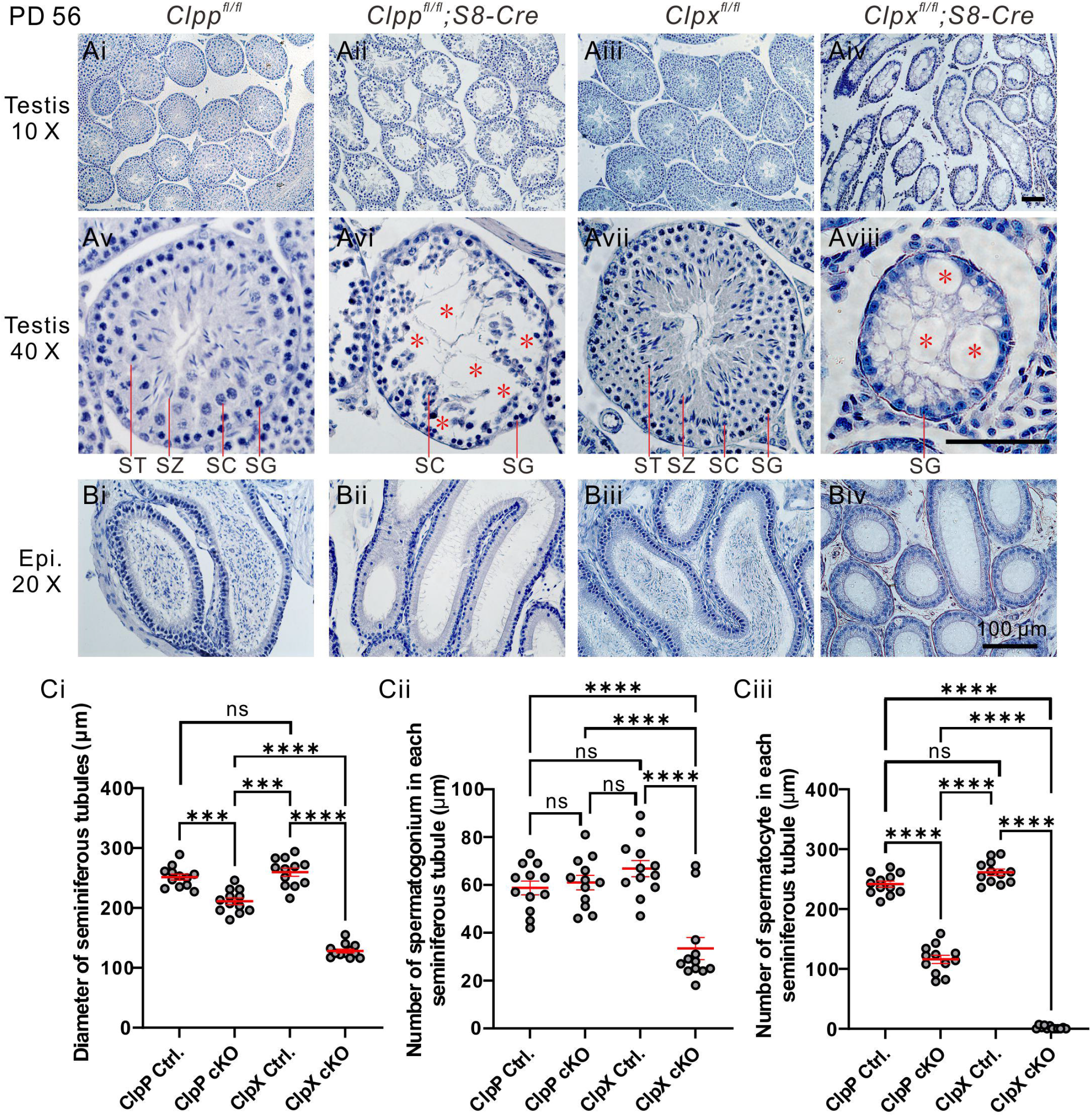
The ClpP/ClpX cKO mouse exhibits abnormal histological structure of testis and disrupted spermatogenesis. (A,B) Testis sections acquired at PD 56 from (Ai,Av) *ClpP*^*fl/fl*^, (Aii,Avi) *ClpP*^*fl/fl*^;Stra8-Cre mice and (Aiii,Avii) *ClpX*^*fl/fl*^, (Aiv,Aviii) *ClpX*^*fl/fl*^*;Stra8-Cre* mice, were stained with hematoxylin. In the *ClpP*^*fl/fl*^*/ClpX*^*fl/fl*^*;Stra8-Cre* testes, the red asterisks indicate abnormal tubules in which the majority of the spermatozoon (SZ), spermatocytes (SC) and post-meiotic spermatids (ST) were absent. SG: spermatogonia. (Bi-Biv) Epididymis sections were also stained with hematoxylin reagent to visualize the histological structure and stored mature sperms. Scale bars are 100 μm. (Ci) Individual dot plot to show the diameter of seminiferous tubules in the ClpP/ClpX control and ClpP/ClpX cKO testes. (Cii) Individual dot plot to show the number of spermatogonia in each seminiferous tumble in the ClpP/ClpX control and ClpP/ClpX cKO testes. (Ciii) Individual dot plot to show the number of spermatocytes in each seminiferous tubles in the ClpP/ClpX control and ClpP/ClpX cKO testes. In (Ci-Ciii), the data represent the mean ± SEM of three independent experiments. ns means no significant difference. ***p < 0.001, ****p < 0.0001.

The ClpX cKO animals exhibited much more severe phenotypes of the seminiferous tubules than the ClpP cKO animals; almost all the germ cells were lost in every tubule under low magnification (Fig. 2Aiv). At high magnification, there were a lot of vacuolar structures in the tubules (red asterisks; Fig. 2Aviii), which were filled with germ cells in the controls (Fig. 2Avii). The spermatogonia in the ClpX cKO remained in the peripheral region of the seminiferous tubules (Fig. 2Aviii). No sperm was observed in the epididymis of the ClpP cKO and the ClpX cKO mice (Fig. 2Bii and Biv), while mature sperms filled up the control epididymis (Fig. 2Bi and Biii).

To quantify the difference in testicular morphology, we determined the diameters of the seminiferous tubules, which were similar between the ClpP control (∼251.5 μm) and the ClpX control (∼259.7 μm). The diameters of the seminiferous tubules in the ClpP cKO (∼211 μm) and the ClpX cKO (∼128 μm) mice were much reduced (Fig. 2Ci). In addition, the number of spermatogonium in the ClpX cKO was significantly lower than the other three groups (Fig. 2Cii), and that the number of spermatocytes (the primary and secondary spermatocytes) in the cKO mice were dramatically reduced when compared with their respective controls (Fig. 2Ciii).

### Deletion of ClpP/ClpX disrupts development and promotes apoptosis of spermatocytes

The ClpP/ClpX cKO testicular sections were immune-stained for γH2A.X (green; Fig. 3A and D) to label DNA double-strand breaks. The sections were co-immuno-stained with SYCP3 (red) for all chromosomes of meiotic cells and the nuclei were stained with DAPI (blue). The expression of γH2A.X in majority of the spermatocytes in the controls (*ClpP*^*fl/fl*^ or *ClpX*^*fl/fl*^ mice; Fig. 3Ai and Di) was concentrated in the sex body, suggesting that these cells were in the pachytene stage or later stages of meiosis. In the ClpP cKO sections, several clusters of spermatocytes in the central zone of the seminiferous tubule retained the γH2A.X signals not only in the sex body but also in the autosomes (abnormal SC, Fig. 3Aiv), suggesting a disruption of meiosis in the ClpP deficient germ cells. Similarly, the ClpX cKO sections exhibited a great number of abnormal spermatocytes at PD 35 with γH2A.X signal retention in the autosomes (yellow arrow heads, Fig. 3Div). The phenotype in the ClpX cKO sections was more severe than that in the ClpP cKO sections.

**Fig. 3.**
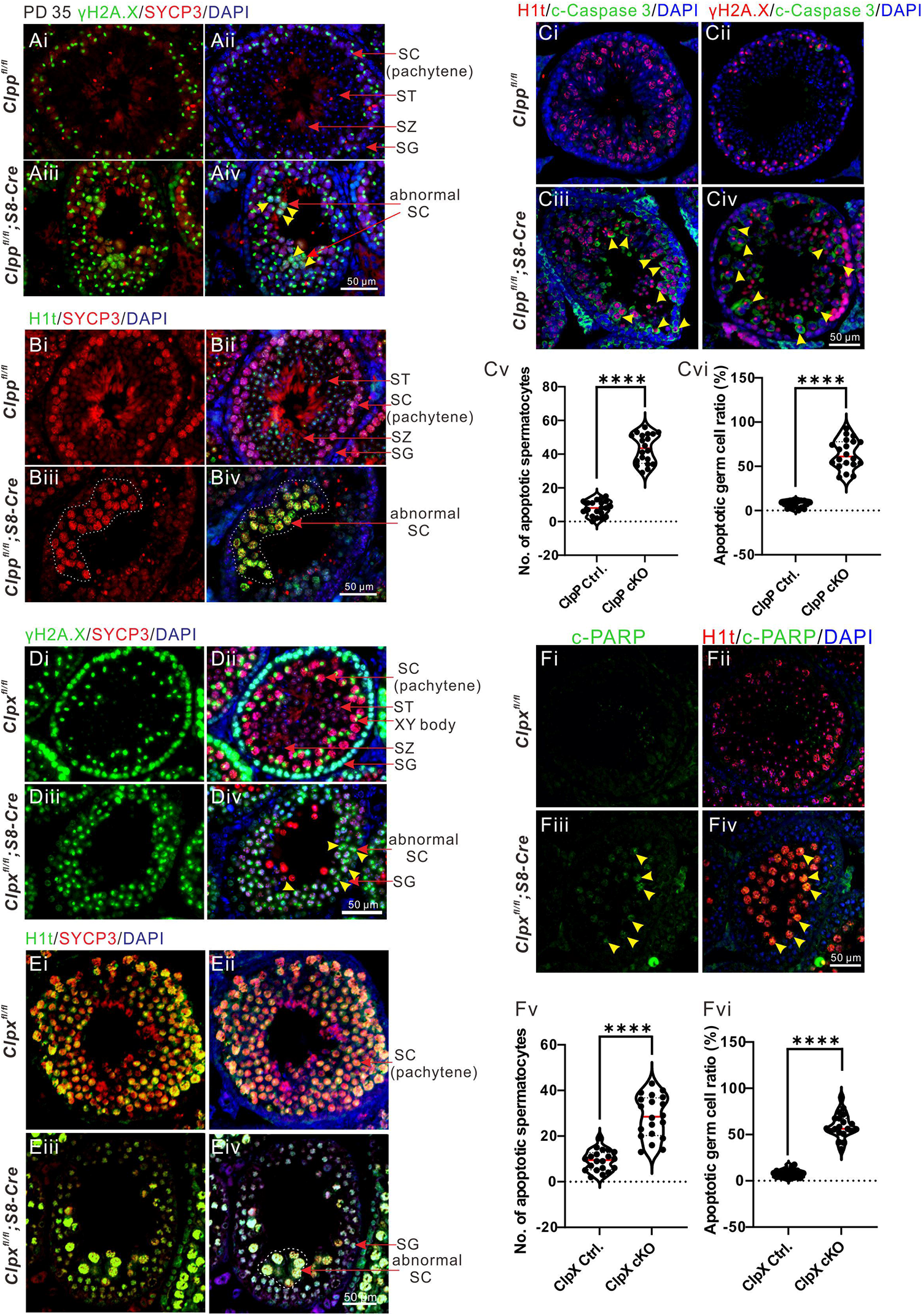
ClpP/ClpX cKO mouse exhibits abnormal spermatocyte differentiation. (A-F) Testis tissue sections from controls and cKO groups were immunolabelled with various primary antibodies (γH2A.X labels double-strand breaks, SYCP3 labels meiotic spermatocytes, H1t labels mid-pachytene spermatocytes, cleaved-Caspase3 or cleaved-PARP labels apoptotic cells and DAPI labels nuclei) and they were co-stained with DAPI (blue). (Cv-Cvi, Fv-Fvi) The individual value plots indicate the quantification of apoptotic spermatocytes and apoptotic germ cell ratio in each section layer of seminiferous tumbles, n=20 for each group. All images were acquired via fluorescent microscope. In (Ciii-Civ, Fiii-Fiv), the yellow arrowheads indicate apoptotic cells. Scale bars are 50 μm. ****: *p*<0.0001.

We further immune-stained the testis sections for H1t, which is normally expressed after the mid-pachytene stage in spermatocytes. The expression pattern of H1t was different between the controls and the ClpP/ClpX cKO mice. More H1t positive cells were observed in the controls than the cKO groups. Majority of the H1t positive cells in the control group were at the late stages of meiosis, including round spermatids and spermatozoa, whereas the H1t positive cells in the ClpP/ClpX cKO mice were bigger in size with pachytene-like appearance and were abnormally aggregated in the seminiferous tubules (Fig. 3B and E). These results suggested that the deletion of ClpP/ClpX disrupted pachytene entry during meiosis.

Since we noticed less germ cells in the ClpP/ClpX cKO testis, we determined apoptosis of germ cells in the mice by immunostaining for apoptotic markers, cleaved-caspase 3 and cleaved-PARP (Fig. 3C and 3F). Compared with the controls, more ClpP/ClpX cKO spermatocytes expressed apoptotic signals (yellow arrow heads; Fig. 3C and F); the number and percentage of apoptotic germ cells were significantly higher in the ClpP/ClpX cKO sections than their respective control (Fig. 3Cv-Cvi and 3Fv-Fvi).

### ClpP or ClpX is required for maintaining meiotic process

We utilized meiotic spreads assay to determine whether the key activities during meiosis were disrupted after knocking out of ClpP or ClpX. The spermatocytes were isolated and divided into 4 meiotic stages, namely leptotene, zygotene, pachytene and diplotene (Fig. 4A). In the control group (*Clpp*^*fl/fl*^ or *Clpx*^*fl/fl*^), the γH2A.X signals were expressed widely in all chromosomes during the leptotene and zygotene stage (Fig. 4Ai-Aii). The signal became concentrated only in the sex chromosome when entry into the pachytene stage (Fig. 4Aiii) and this specific expression pattern was retained through the diplotene stage (Fig. 4Aiv). We noticed that the γH2A.X signal could not be concentrated in the sex body and remained at a pachytene-like stage in both cKO mice (Fig. 4 Avii and Axi). Moreover, the strong retention of the γH2A.X expression was also found in the autosomes of the diplotene-like stage (Fig. 4Aviii and Axii). The percentages of cells at different meiotic stages in the cKO and control mice were compared. Majority of the ClpP/ClpX cKO spermatocytes stuck during transformation from the zygotene to the pachytene stage (Fig. 4B).

**Fig. 4.**
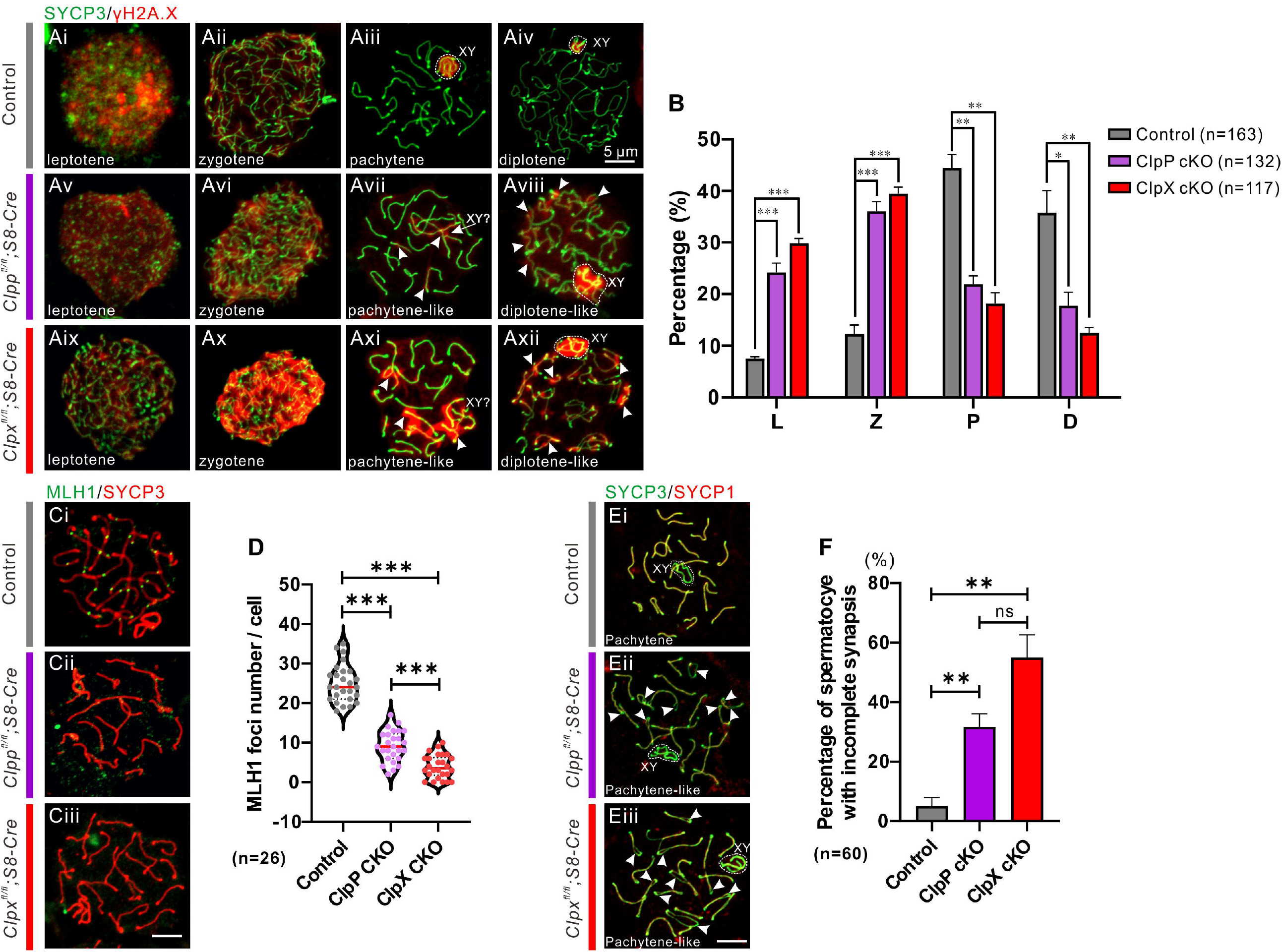
Comparison of meiotic chromosome spreads from controls, ClpP cKO and ClpX cKO spermatocytes. (A,C,E) Harsh chromosome spreads were prepared before immunolabeling procedures were conducted with the γH2A.X, MLH1, SYCP1 and SYCP3 antibodies. The SYCP3 antibody was used to label meiotic chromosomes. (A) Spermatocytes are shown at different stages. (B) Bar chart shows the mean ± SEM of percentage of spermatocytes at these stages (i.e., L: leptotene, Z: zygotene, P: pachytene, and D: diplotene) were quantified. n=163 for control spermatocytes, n=132 for ClpP cKO spermatocytes and n=117 for ClpX cKO spermatocytes. (C) Spermatocytes in different groups were immunolabelled with the anti-MLH1 (in green) and anti-SYCP3 (in red) antibodies. (D) The number of MLH1 foci were quantified in the controls, ClpP cKO and ClpX cKO spermatocytes (n=26 for each group). (E) Spermatocytes in different groups were immunolabelled with the anti-SYCP1 (in red) and anti-SYCP3 (in green) antibodies. The dashed line cycled XY body and white arrow heads indicate chromosomes asynapsis. (F) Bar chart shows the mean ± SEM of percentage of spermatocytes with incomplete synapsis (n=60 for each group). Scale bars are 5 μm. *: *p*<0.05; **: *p*<0.01; ***: *p*<0.001; ns: non-significant difference.

We next immune-stained the spermatocytes for MLH1 to check whether crossover was affected. In the control pachytene spermatocytes, the chromosomes were positively stained with the anti-SYCP3 antibody (red), and the MLH1 foci were expressed on each chromosome (green, Fig. 4Ci), illustrating normal crossover activities during meiosis. However, almost no MLH1 foci was detected on the chromosomes in the pachytene-like stages of the ClpX cKO spermatocytes (Fig. 4Ciii). The number of MLH1 foci in the ClpP cKO spermatocytes was remarkably decreased (Fig. 4Cii). We further quantified the number of MLH1 foci in each cell. The data showed significant decrease in the ClpP/ClpX cKO cells relative to the controls, and that the decrease was more severe in the ClpX cKO cells than the ClpP cKO cells (Fig. 4D).

Next, we tested whether the chromosomal synapsis was affected in the ClpP/ClpX deleted spermatocytes. The meiotic spreads assays with immune-staining for a synapsis specific marker, SYCP1 were performed (Fig. 4E). Expression of SYCP1 on each autosome rather than the sex chromosome was observed in the pachytene spermatocytes of the control mice, indicating the occurrence of normal chromosome synapsis (Fig. 4Ei). However, the ClpP/ClpX cKO spermatocytes exhibited asynapsis of the autosomes at the pachytene-like stage; the sister chromosomes were separated (white arrow heads) and the expression of SYCP1 (red) was absent (Fig. 4Eii and Eiii). Quantification of the percentages of incomplete synapsis at the pachytene/pachytene-like stage showed that 5% of the control pachytene spermatocytes presented asynaptic chromosomes, whereas the percentage increased to 31.6% and 55% in the ClpP cKO and the ClpX cKO spermatocytes, respectively (Fig. 4F). Our data demonstrated the the ClpP and ClpX were required for spermatocyte chromosomal synapsis at the pachytene stage, as well as the following crossover activities. The lack of ClpP or ClpX expression in the spermatocytes attenuated the meiotic process, specifically during the zygotene-to-pachytene transition.

### ClpP/ClpX is crucial for telomere-nuclear envelope attachment and α-tubulin formation

To further understand the role of ClpP/ClpX in meiosis, we investigated the dynamic of telomeres during meiosis. Mild meiotic spreads assay was performed with immunostaining for telomeres and nuclear envelope using the anti-TRF1 antibody and the anti-Lamin B1 antibody, respectively (Fig. 5A). Images of a mid-layer of the control spermatocytes demonstrated nice attachment of the telomeres (TRF1 foci in green) to the nuclear envelope (Lamin B1 labeled membrane in blue), reflecting a normal telomeres-nuclear envelope attachment during meiosis (Fig. 5Ai-Aii). On the other hand, the ClpP/ClpX cKO cells showed remarkable number of telomeres that failed to attach to the nuclear envelope and located in the central region of the nucleus (white asterisks; Fig. 5Aiii-Avi). We counted the number of detached TRF1 foci and found significant increase in the number in the ClpP/ClpX cKO cells when compared with the controls (Fig. 5Bii). The ratio of cells with failure of telomere-nuclear envelope attachment to the total cells was also significantly increased in the ClpP/ClpX cKO spermatocytes (Fig. 5Bi).

**Fig. 5.**
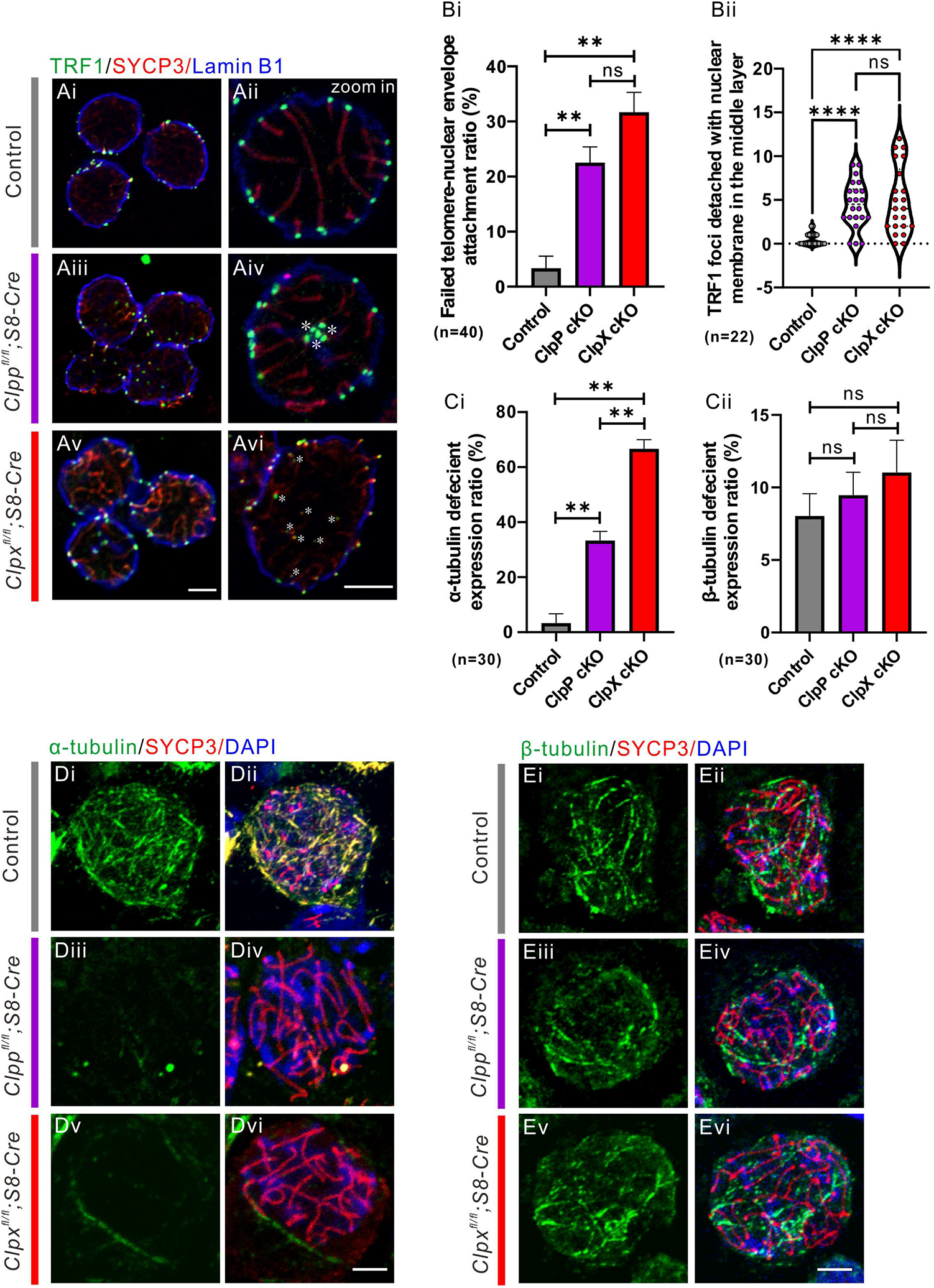
Evaluation of the attachment between telomere and NE during meiosis and tubulin formation in ClpP/ClpX cKO spermatocytes. (A) The spermatocytes from control, ClpP cKO and ClpX cKO groups were prepared to perform mild chromosome spreads before immunolabelling with an anti-TRF1 (in green), anti-SYCP3 (in red) and anti-Lamin B1 (in blue) antibodies, Lamin B1 acts as a marker to label nuclear membrane. White asterisks indicate detached telomeres with NE. (B) Quantification of (Bi) failed telomere-NE attachment ratio in spermatocytes in each group (n=40, bar chart shows the mean ± SEM) and (Bii) the number of TRF1 foci which detached with NE in the middle layer (n=22 spermatocytes for each group). (D-E) The spermatocytes with cytospin were immunolabeled with (D) anti-alpha-tubulin and (E) anti-beta-tubulin antibodies to visualize the formation of tubulin-based cytoskeleton in control, ClpP cKO and ClpX cKO spermatocytes. These samples were co-labeled with SYCP3 to confirm the mitotic germ cells. (C) Bar charts show the mean ± SEM of (Ci) alpha-tubulin or (Cii) beta-tubulin deficient spermatocyte ratio (n=30 for each group). All images were captured by confocal microscopy. Scale bars are 5 μm. **: p<0.01; ****: p<0.0001; ns: non-significant difference.

During the experiments, we found that the ClpP/ClpX cKO spermatocytes were fragile when compared with the control spermatocytes. It is known that chromosome and telomere dynamics are associated with cytoskeleton functions (*29, 30*). Therefore, the cytoskeleton of the ClpP/ClpX cKO cells was checked. The spermatocytes were purified and cytospun onto a glass slide. Immunostaining for α-tubulin and β-tubulin showed that there was no difference in β-tubulin expression intensity when compared with the controls, but the α-tubulin formation was severely attenuated in the ClpP/ClpX cKO spermatocytes (Fig. 5D-E). Attenuation of α-tubulin formation was found in more than 60% of the ClpX cKO spermatocytes and around 35% of the ClpP cKO spermatocytes; both percentages were significantly higher than that in the control spermatocytes (Fig. 5Ci). No significant difference was found for β-tubulin formation among these groups (Fig. 5Cii).

### Deletion of ClpP/ClpX affects mitochondrial quantity and morphology

We performed electron microscopy (EM) to study the mitochondria in the germ cells. In the control cells, the mitochondrial morphology in the spermatogonia (SG, Fig. 6Ai-Aiv) and spermatocytes (SC, Fig. 6Ai-Aiv) was normal with elongated shape and well-formed inner/outer membranes, intermembrane space and cristae. The mitochondrial morphology in the corresponding ClpP/ClpX cKO cells were different. Almost no elongated mitochondria could be detected in the ClpP/ClpX cKO spermatocytes. In addition, the cristae structure was disrupted and vacuolar structures were observed in the mitochondria (Fig. 6Av-Axii). The quantification data shows no statistical difference in the number of mitochondria and the mitochondrial length-width ratio between the controls and the ClpP/ClpX cKO spermatogonia (Fig. 6Bi-Bii). However, these parameters were significantly lower in the ClpP and ClpX deficient spermatocytes than the controls (Fig. 6Bi and Biii). The WT spermatocytes harbored more mitochondria than the spermatogonia (Fig. 6Bi).

**Fig. 6.**
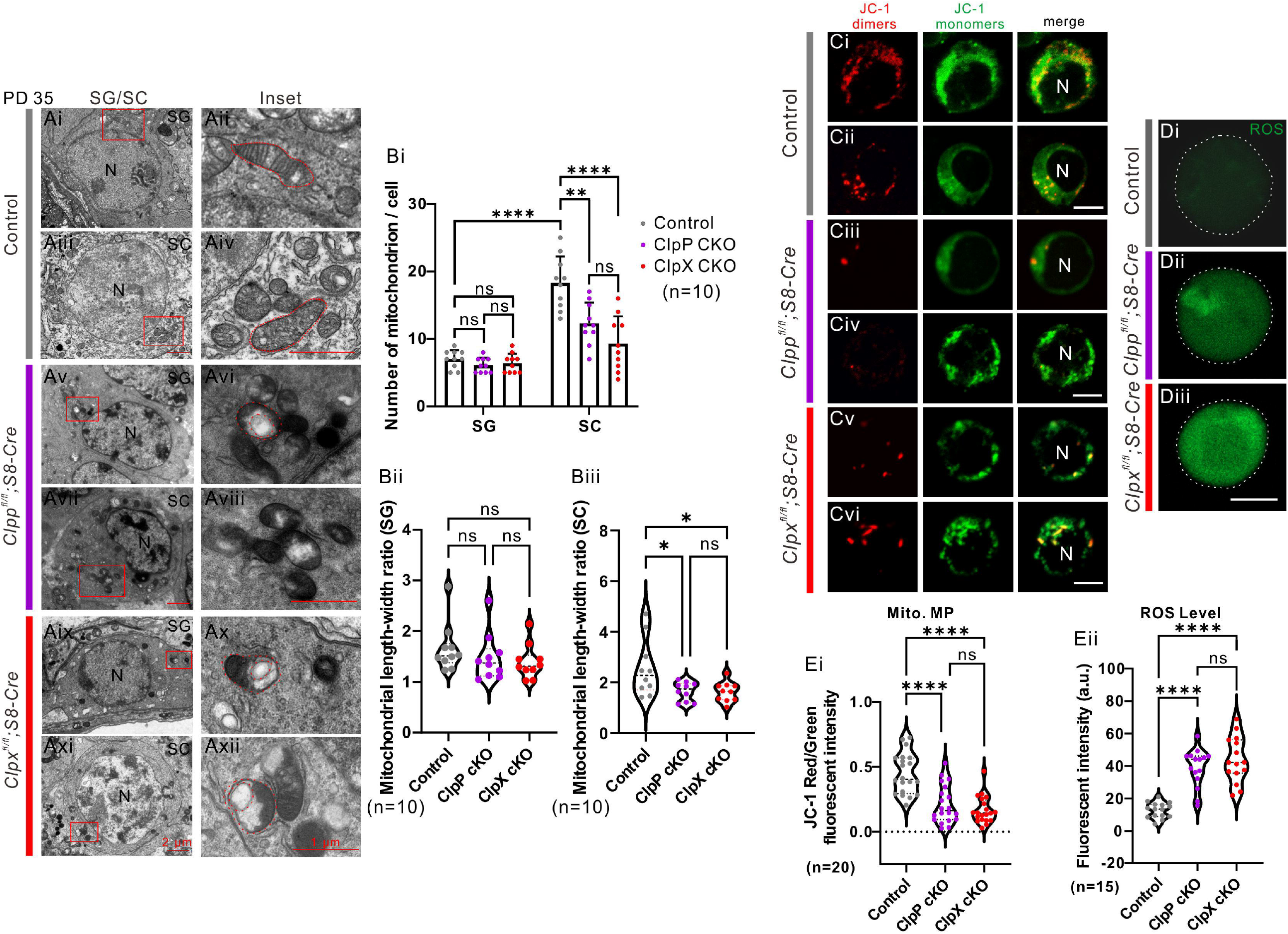
Evaluation of mitochondrial expression and functions in germ cells. (A) The testis tissue sections were prepared for transmission electro-microscopy (TEM). Spermatogonium (SG) and spermatocytes (SC) were captured to visualize the morphology of mitochondria. N means nucleus. Scale bars are 2 μm in the SG/SC panel and 1 μm in the inset panel, respectively, as labeled in the figure. (Bi) Number of mitochondria in each cell captured layer were quantified in both SG and SC cells. Mitochondrial length/width ratio was also quantified in (Bii) SG and (Biii) SC, respectively. (C) The mitochondrial membrane potential was measured via JC-1 staining in the (Ci-Cii) control, (Ciii-Civ) ClpP cKO and (Cv-Cvi) ClpX cKO spermatocytes, JC-1 aggregates were labelled in red and JC-1 monomers were labelled in green, N means nucleus, scale bars are 10 μm. (D) The level of ROS was visualized in these three groups via labeling with carboxy-H_2_DCFDA (in green), scale bar is 10 μm. Images in Ci-Diii were captured via confocal microscope. (Ei) Mitochondrial membrane potential was quantified via the ratio of red/green fluorescent intensity, 20 spermatocytes were quantified in each group. (Eii) The ROS level of spermatocytes in three groups was quantified via fluorescent intensity, n=15 for each group. *: p<0.05, **: p<0.01; ****: p<0.0001; ns: non-significant difference.

### Deletion of ClpP or ClpX affects mitochondrial function in spermatocytes

Next, we studied the effects of ClpP or ClpX deficiency on mitochondrial functions. First, JC-1 staining in isolated live spermatocytes was conducted. Red fluorescence reflects JC-1 aggregates and green fluorescence reflects JC-1 monomers, and the ratio of red/green fluorescence is a measure of the mitochondrial membrane potential. The control spermatocytes exhibited stronger red fluorescent signals than the ClpP/ClpX cKO spermatocytes, whereas the green fluorescent signals were comparable among them (Fig. 6C). The red/green fluorescence ratios were significantly lower in the ClpP/ClpX cKO spermatocytes than the controls (Fig. 6Ei), suggesting that the mitochondrial membrane potential was attenuated in the ClpP/ClpX deficient spermatocytes.

We also evaluated the ability of ROS level recovery and found that the ClpP/ClpX deficient spermatocytes could not reduce the ROS level back to the normal value after H_2_O_2_ pretreatment; both groups showed a significant higher ROS level relative to the controls (Fig. 6Eii). In addition, quantitative PCR assays showed a significant altered expression of several respiratory chain complex genes including Atp5a1, *Ndufv1, Cox1, Uqcrc2* and *Sdhb* in the ClpP/ClpX cKO spermatocytes (fig. S2). Based on these observations, we concluded that the deficiency of ClpP or ClpX attenuated the mitochondrial functions in spermatocytes.

### Transcriptome profiling and m6A-seq analysis of ClpP/ClpX cKO spermatocytes

The germ cells from PD 35 mice were isolated and purified by gravity sedimentation. We selected the pachytene and the pachytene-like spermatocytes (diameter 18-25 μm). RNA sequencing and m6A sequencing were performed on the spermatocytes from the controls, ClpP cKO and ClpX cKO mice (three mice for each group). Principal components analysis (PCA) showed clustering of the 3 groups of samples except the sample WT Ctrl. 3, which was away from the other two control samples WT Ctrl. 1 and 2 (Fig. 7A). When compared with the controls, the ClpP cKO spermatocytes had 1047 down-regulated genes and 147 up-regulated genes, while the ClpX cKO spermatocytes showed 3175 down-regulated genes and 1496 up-regulated genes (fig. S3). Gene ontology (GO) enrichment analysis confirmed our observed phenotypes and highlighted changes in spermatogenesis, spermatid differentiation and development in the ClpP cKO and ClpX cKO samples (Fig. 7B-C). The analysis further suggested that the ion channel activity, DNA packaging complex and nucleosome were affected (Fig. 7B-C). Heatmap of the transcriptome profiles of the 9 samples showed clustering of the WT controls, while the profile of the ClpX cKO 1 was mixed with that the ClpP cKO profiles (Fig. 7D). Among the differential expressed genes between the ClpP cKO and the WT and between the ClpX and the WT, 1005 of them were common in the two comparisons (Fig. 7E). GO analysis showed that these genes were enriched for the process of sperm formation and motility, ion transport, aminoacyl-tRNA biosynthesis, and ADP metabolism. (Fig. 7F).

**Fig. 7.**
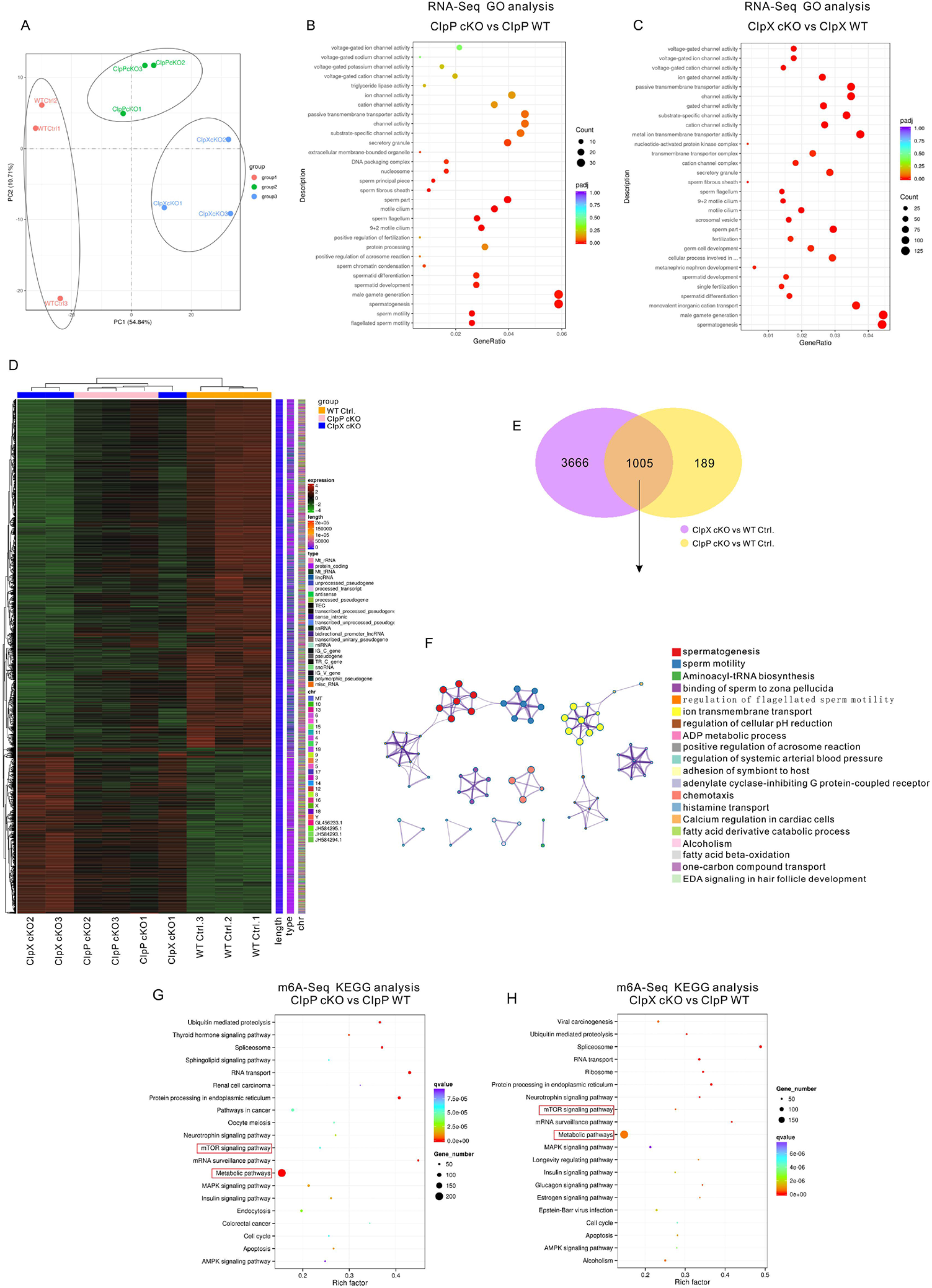
Transcriptome profiling and m6A sequencing analysis of spermatocytes in WT control, ClpP cKO and ClpX cKO groups. (A) PCA plot of three groups of spermatocytes were shown after RNA sequencing from n=3 mice in each group. The dot plots of GO analysis were shown for (B) ClpP Cko and (C) ClpX cKO when compared with WT controls. (D) Heatmap and cluster analysis of the overview of gene expression patterns in all 9 samples from WT control, ClpP cKO and ClpX cKO groups. (E) Venn analysis shows common changed genes after deletion of ClpP and ClpX in spermatocytes. (F) Metascape network plots showing the relationship of the enriched GO terms in 1005 common genes from Venn analysis. (G-H) Spermatocytes were purified from PD 35 WT control or ClpP/ClpX cKO mice testis and sent for m6A sequencing. Dot plots of KEGG analysis showed gene enrichment in related pathways after knocking out of ClpP or ClpX.

We also performed m6A-seq of the ClpP/ClpX cKO spermatocytes to evaluate the change in RNA methylation level. Based on Kyoto Encyclopedia of Genes and Genomes (KEGG) analysis, we found majority of the highlighted genes were enriched in metabolic pathways and both the ClpP cKO and ClpX cKO spermatocytes showed alteration of RNA methylation in mTOR signaling related genes (Fig. 7G-H). The conclusion was supported by the expression levels of the mTOR signaling genes in the controls, ClpP cKO and ClpX cKO groups (fig. S3E). Based on the heatmap results, majority of the mTOR signaling genes, including *Akt1, Rps6, Mapk1, Rhoa*, etc., were up-regulated, whereas around one-third of the mTOR signaling genes, including *Braf, Atg13, Sspo* etc., were down-regulated after deletion of ClpP/ClpX (fig. S3E)

### mTORC1 signaling is over-activated in ClpP/ClpX deleted spermatocytes

We confirmed over-activation of the mTORC1 signaling pathway in the ClpP/ClpX cKO spermatocytes by western blotting (fig. S1). Next, we conducted *in vivo* pharmacological treatment with a widely known mTORC1 inhibitor, rapamycin. The ClpP/ClpX cKO mice were intraperitoneally injected with PBS (control) or rapamycin (8 mg/kg) every other day from PD 14 and the mice were sacrificed on PD 35 (Fig. 8A). Histological analysis of testis and epididymis tissue failed to observe the round spermatids or spermatozoon, respectively in the ClpP/ClpX cKO mice receiving PBS. Instead, vacuolar structures (red asterisks) probably formed after germ cell loss were observed (Fig. 8Bi and Bv). Surprisingly we found both round spermatids and elongated spermatozoa in the testicular sections of the rapamycin injected ClpP/ClpX cKO mice at PD 35 (Fig. 8Bii-Biii and Bvi-Bvii, fig. S4). The observation indicated that the inhibition of mTORC1 signaling partially rescue the meiotic process of germ cells in the ClpP/ClpX cKO mice, though mature sperms could still not be found in their epididymis (Fig. 8Biv and Bviii). Quantification of the histological examination shows a significant increase in the percentage of seminiferous tubules with elongated spermatozoa in the rapamycin treated ClpP/ClpX cKO mice (Fig. 8Bix). There was no further improvement in the rescue outcome when the rapamycin treatment was extended to PD 56; no mature sperm could be detected in either the testis or the epididymis (fig. S4). The sizes of the rapamycin treated cKO testes remained comparable with the PBS injected ClpP/ClpX cKO testes at PD 56 (fig. S4C). In addition, we found that the seminiferous tubules of the rapamycin-treated tissues were more closely packed than that of the controls (fig. S4C, top panel).

**Fig. 8.**
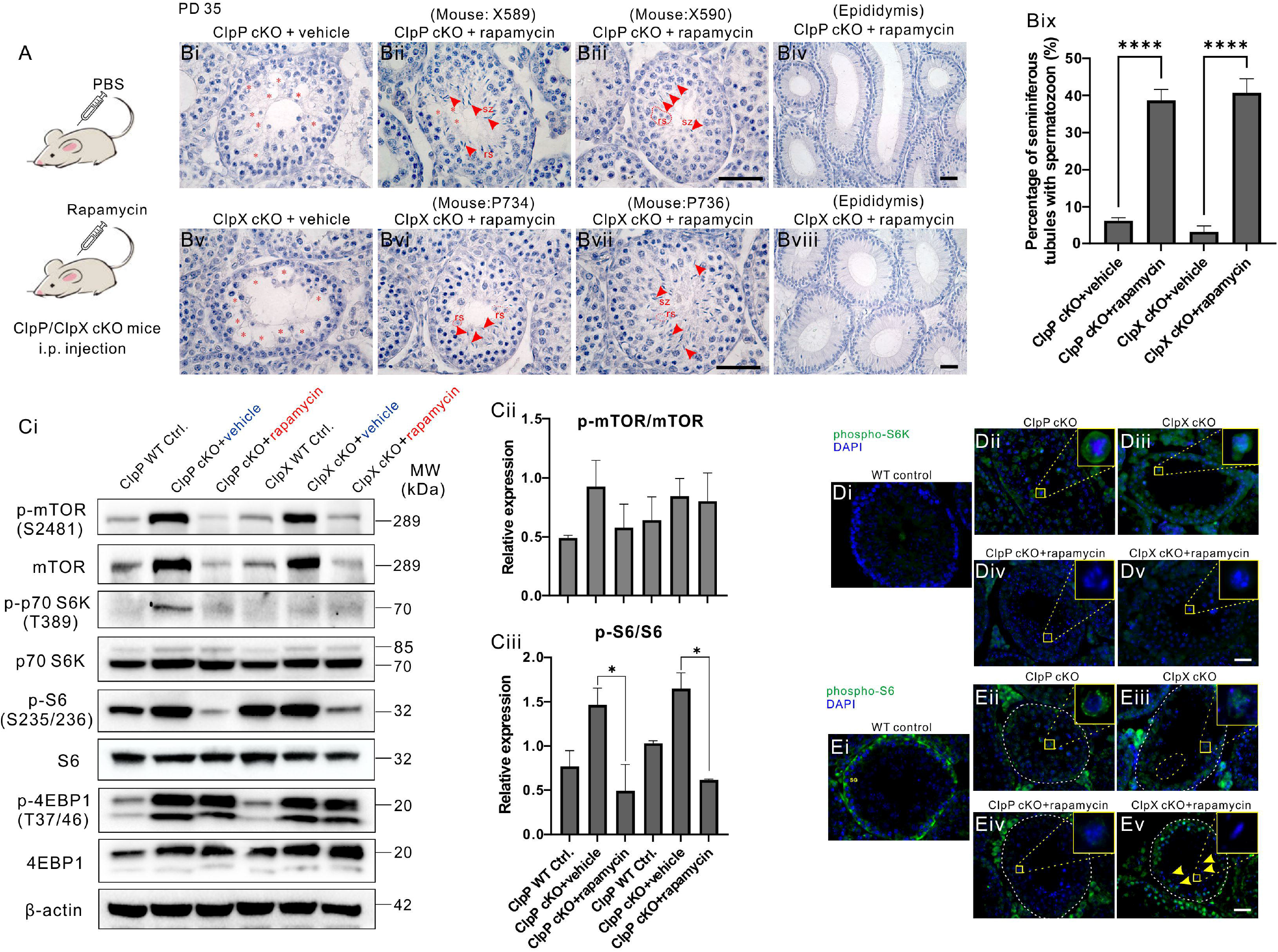
In vivo pharmacological treatment of rapamycin to inhibit mTORC1 in ClpP/ClpX cKO mice. (A) Carton diagram to describe the plan of drug injection. (B) Histological studies showed PD 35 testis and epididymis sections with hematoxylin staining. X589 and X590 are mice ear tags for ClpP cKO injected with rapamycin, P734 and P736 are mice ear tags for ClpX cKO injected with rapamycin. Red asterisks labeled empty regions with germ cell died out, red arrow heads indicated elongated spermatozoon (sz) and red dashed line circled round spermatids (rs). (Bix) The percentage of spermatozoon harbored seminiferous tubules was quantified. (Ci) The protein expression level of mTOR signaling pathways was shown via western blot analysis, the phosphorylation level of (Cii) mTOR and (Ciii) S6 were quantified. Immunolabeling of (D)phosphor-S6K (in green) and (E) phosphor-S6 (in green) in ClpP/ClpX cKO or ClpP/ClpX cKO injected with rapamycin testis tissue sections, DAPI (in blue) was used to label nucleus. Images were captured via fluorescent microscope. Scale bars are 50 μm. *: *p*<0.05; ****: *p*<0.0001.

To confirm the suppression of mTORC1 after rapamycin treatment, we performed western blot analysis on the purified spermatocytes. The phospho-mTOR level and pan-mTOR were up-regulated in the ClpP cKO and the ClpX cKO spermatocytes, and they returned to level found in the WT cells after rapamycin treatment (Fig. 8Ci-Cii). The two main substrates of mTORC1, p70-S6K and S6, were also evaluated. The expression of phospho-p70 S6K was almost undetectable in the ClpP/ClpX WT controls. On the other hand, we detected a remarkable increase in phospho-p70 S6K expression in the ClpP cKO injected with PBS, consistent with the activation of p70 S6K in a ClpP deficient condition (Fig. 8Ci). The activation was inhibited after rapamycin treatment and the band was much weaker compared with the ClpP cKO group (Fig. 8Ci). A very weak band was seen in the ClpX cKO+vehicle and the ClpX cKO+rapamycin groups. The pan-p70 S6K levels were comparable among these groups, suggesting that the phosphorylation but not the total amount of p70 S6K was increased in the absence of ClpP or ClpX (Fig. 8Ci). The pan-S6 level was slightly decreased in the ClpP/ClpX cKO spermatocytes when compared with the WT controls, while the phospho-S6 in ClpP/ClpX cKO groups showed dramatic increase when compared with ClpP/ClpX WT controls. In the rapamycin treated ClpP/ClpX cKO groups, the phospho-S6 levels were significantly inhibited (Fig. 8Ci and Ciii). Another substrate of mTORC1, 4EBP, was also studied. Both the phospho-4EBP1 and pan-4EBP1 showed dramatic increase in the ClpP/ClpX cKO cells relative to the ClpP/ClpX WT controls, and rapamycin treatment could not decrease the expression of phospho-4EBP1 in the ClpP/ClpX cKO cells, consistent with a previous study (*31*). The expression of β-actin showed was comparable in all the groups (Fig. 8Ci).

Since the expression of phospho-S6K is relatively weak and cannot be accurately compared by western blot analysis, we evaluated the expression pattern of phospho-S6K/S6 in testicular sections by immunostaining. In the WT controls, almost no obvious phospho-S6K could be detected in the spermatocytes (Fig. 8Di), while the ClpP/ClpX cKO spermatocytes showed strong cytoplasmic expression of phospho-S6K (green; Fig. 8Dii-Diii). The expression of phospho-S6K in the ClpP/ClpX cKO spermatocytes disappeared after rapamycin treatment (Fig. 8Div-Dv). Interestingly, we could also see strong phospho-S6 signals in the spermatogonia but not the spermatocytes of ClpP/ClpX WT controls (Fig. 8Ei), whereas the expression of phospho-S6 decreased in the spermatogonia but increased in the spermatocytes of the ClpP and ClpX cKO testis (Fig. 8Eii-Eiii). We also found some fragmented nuclei in the ClpX cKO seminiferous tubule (circled in yellow dotted line, Fig. 8Eiii), suggested apoptosis of the germ cells in the ClpX deficient testes. The phenotype could be partially rescued after rapamycin treatment, with the expression of phospho-S6 in the spermatocytes decreased to levels comparable to the controls and partial recovery of the phospho-S6 expression in the spermatogonia after the treatment. Some elongated spermatozoa were seen (yellow arrow heads, Fig. 8Eiv-Ev).

## DISCUSSION

The mitochondrial function is widely recognized as one of the key factors for germ cell development (*32-34*). However, the role of ClpP/ClpX in meiosis and spermatogenesis is not fully understood. Although some recent studies reported about ClpP-mediated mitochondrial quality control in ovary aging (*22, 23*), the research of ClpP/ClpX-mediated mitochondrial quality control in male germ cell development and spermatogenesis is still rare. In this study, we utilized Cre-LoxP and *Stra8*-driven Cre system (*35*) to generate male germ cell specific knockout of ClpP or ClpX in mice. The deficiency of ClpP or ClpX severely affected mitochondrial functions in spermatocytes, which further disrupted meiotic events and progression, and eventually failure of spermatogenesis.

We found genetic compensation of the two genes in the ClpP and the ClpX knockout animals. Compared with controls, the ClpP cKO spermatocytes expressed more ClpX protein, and vice versa for the ClpX cKO spermatocytes. The observation is in line with a previous study showing remarkable ClpX accumulation in many organs, including testis and ovary of ClpP deleted mice (*25*). The ClpP/ClpX cKO testes were smaller than the controls. Testicular histology revealed that the decreased testis size was due to the loss of spermatocytes. Deletion of ClpP and ClpX affected mainly the spermatocytes and did not have much effect on the spermatogonia. This differs from deletion of mitofusin 2 protein, which not only leads to apoptosis of the spermatocytes but also affects survival of the spermatogonia (*36*).

In this study, we focused on the meiosis process as the loss of spermatocytes could be related to disruption of meiosis. Based on the expression pattern of γH2A.X (DNA double-strand break marker) and H1t (mid-pachytene spermatocyte marker), we found that the ClpP or ClpX deficient spermatocytes could not proceed to the pachytene stage successfully. The observation is in line with a previous study showing failure of progression of spermatogenesis especially at the zygotene-pachytene transformation stage in germ cells of *Mfn1 and Mfn2* mutant mice (*37*). Here, our meiotic spreads assay showed that majority of the control cells were at the pachytene or diplotene stages while most of the ClpP/ClpX cKO cells were stuck at the leptotene or the zygotene stages on PD 35. In addition, our immunostaining of MLH1 and SYCP1 on the meiotic spreads demonstrated that ClpP and ClpX were required for chromosome synapsis. It is known that crossover will not happen if the chromosomes are in a condition of asynapsis (*38*). We further showed that the meiotic obstruction eventually led to apoptosis of the cKO spermatocytes, which provided an explanation of the abnormal testicle morphology in the mutant mice. Based on these data, we conclude that ClpP and ClpX are required for the maintenance of spermatocytes, but not the spermatogonia.

The most direct role of mitochondria in meiosis is energy supply, as the various activities of meiosis, such as chromosomal dynamics, require ATP (*39*). In meiotic prophase, the telomeres rather than the centromere assemble the chromosome and bind to the nuclear envelope (*40, 41*). Deficiency of ClpP or ClpX induced detachment of the telomere from the nuclear envelope; the single layer of high-resolution confocal images clearly demonstrated the dropped telomeres (TRF1 labeled) in the central region of the nuclei instead of attachment of the telomeres to the nuclear envelope as in the controls. Similar phenomenon was observed in the MFN2 cKO spermatocytes (*34*). Therefore, we studied the formation of microtubule in the ClpP/ClpX deleted spermatocytes, as cytoskeleton is known to mediate chromosome dynamics (*42*). Surprisingly, we found that ClpP/ClpX deletion disrupted formation of α-tubulin but not β-tubulin. Electron microscopy showed abnormal mitochondrial morphology in the ClpP/ClpX cKO germ cells. The physiological functions of mitochondria were also disrupted in the ClpP/ClpX cKO spermatocytes as demonstrated by two typical criteria of mitochondrial dysfunction, namely, decrease in mitochondrial membrane potential and increase in ROS level. These data clearly demonstrated that the deficiency of ClpP/ClpX impaired mitochondrial functions in germ cells, resulting in disruption of meiosis.

Both the transcriptome analysis and the m6A-sequencing indicate significant alteration of metabolic pathways in the mutant spermatocytes. We determined the expression of mTOR signaling and surprisingly found mTORC1 signaling pathway was activated in the ClpP/ClpX deleted spermatocytes. Therefore, we tested whether manipulation of the mTORC1 activity by rapamycin could rescue the effect of ClpP/ClpX deficiency in spermatogenesis and meiosis. Our *in vivo* rapamycin administration experiment demonstrated the treatment down-regulated mTORC1 signals with decreased phosphorylation of two main substrates of mTORC1, S6K and S6. The increase in the phosphorylation level of 4EBP1 observed after rapamycin injection was consistent with a previous study (*31*). These molecular changes contributed to phenotypic changes in the testicular tissue with reappearance of round spermatids and spermatozoa in the rapamycin treated animals. The observations indicate partial rescue of the meiotic process after manipulation of the metabolism in the germ cells. Unfortunately, we could not observe any mature sperm in the epididymis of the treated mice, this might be because signaling pathway(s) other than mTORC1 were altered and affected meiosis in the ClpP/ClpX mutated male germ cells. Thus, rapamycin treatment could not totally rescue all the affected pathways. Indeed, our sequencing data revealed alterations of many other signaling and biological processes upon ClpP/ClpX deficiency. These data support the conclusion that the mTORC1 signaling is over-activated in the ClpP/ClpX deficient spermatocytes.

In summary, our study demonstrated the role of ClpP and ClpX, in spermatogenesis. We showed that they are essential to maintain normal mitochondrial morphology and functions in the male germ cells, and their deletion in the germ cells disrupts meiosis, which eventually leads to apoptosis of spermatocytes and decrease in testicular size. The ClpP or ClpX deficient mice are infertile and lack mature sperms in the testis and epididymis. High-throughput sequencing analysis highlighted alteration of metabolic pathways in the mutant spermatocytes. We confirmed dramatic upregulation of the mTOR signaling in the ClpP/ClpX cKO spermatocytes. Rapamycin *in vivo* treatment down-regulated the mTORC1 signaling and partially rescued the testicular phenotype of the mutant mice. Our results provide a novel insight of ClpP/ClpX complex in mTOR-mediated metabolism during meiosis and spermatogenesis in mice.

## Acknowledgments

Authors would like to express special thanks to Dr. Kui Liu from The University of Hong Kong for his suggestions and guidance in germ cell isolation and meiosis studies. Thanks also go to Dr. Guangxin Li from Peking University Shenzhen Hospital for his kind discussion on the work of mTOR signaling pathways.

## Funding

This work was supported by:

National Natural Science Foundation of China (No. 81971453);

Shenzhen Fundamental Research Program, China (No. JCYJ20200109150429414);

Shenzhen Science and Technology Program, China (No. RCYX20200714114705073, No. KQTD20190929172749226).

## Author contributions

Conceptualization: C.G., T.W.

Methodology: C.G., T.W.

Investigation: C.G., X.Y., J.G., J.Zheng, R.H.

Animal breeding: C.G., X.Y., J.G., Z. Hu, R.H., Z. Hai, J.S.

Funding acquisition: W.Y., J.Zhang, T.W.

Project administration: C.G., T.W.

Supervision: C.G., T.W.

Writing – original draft: C.G.

Writing – review & editing: C.G., W.Y., T.W.

## Data and materials availability

All data are available in the main text or the supplementary materials. The datasets presented in this study can be found in online repositories. The names of the repository/repositories and accession number(s) can be found below: https://www.ncbi.nlm.nih.gov/, GSE220600.

## REFERENCE

1. A. M. Van Der Bliek, M. M. Sedensky, P. G. Morgan, Cell Biology of the Mitochondrion. Genetics 207, 843–871 (2017).

2. M. Bonora, S. Patergnani, A. Rimessi, E. De Marchi, J. M. Suski, A. Bononi, C. Giorgi, S. Marchi, S. Missiroli, F. Poletti, M. R. Wieckowski, P. Pinton, ATP synthesis and storage. Purinergic Signalling 8, 343–357 (2012).

3. R. C. Scarpulla, Transcriptional Paradigms in Mammalian Mitochondrial Biogenesis and Function. Physiological Reviews 88, 611–638 (2008).

4. J. St-Pierre, I. Topisirovic, Nucleus to Mitochondria: Lost in Transcription, Found in Translation. Developmental Cell 37, 490–492 (2016).

5. A. Ferrer-Vaquer, M. Barragán, A. Rodríguez, R. Vassena, Altered cytoplasmic maturation in rescued in vitro matured oocytes. Human Reproduction 34, 1095–1105 (2019).

6. A. S. Moore, S. M. Coscia, C. L. Simpson, F. E. Ortega, E. C. Wait, J. M. Heddleston, J. J. Nirschl, C. J. Obara, P. Guedes-Dias, C. A. Boecker, T.-L. Chew, J. A. Theriot, J. Lippincott-Schwartz, E. L. F. Holzbaur, Actin cables and comet tails organize mitochondrial networks in mitosis. Nature 591, 659–664 (2021).

7. E. Kummer, N. Ban, Mechanisms and regulation of protein synthesis in mitochondria. Nature Reviews Molecular Cell Biology 22, 307–325 (2021).

8. N. U. Naresh, C. M. Haynes, Signaling and Regulation of the Mitochondrial Unfolded Protein Response. Cold Spring Harbor Perspectives in Biology 11, a033944 (2019).

9. H. Weidberg, A. Amon, MitoCPR—A surveillance pathway that protects mitochondria in response to protein import stress. Science 360, eaan4146 (2018).

10. M. J. Freitas, S. Vijayaraghavan, M. Fardilha, Signaling mechanisms in mammalian sperm motility†. Biology of Reproduction 96, 2–12 (2017).

11. M. Boguenet, P.-E. Bouet, A. Spiers, P. Reynier, P. May-Panloup, Mitochondria: their role in spermatozoa and in male infertility. Human Reproduction Update 27, 697–719 (2021).

12. K. Nakada, A. Sato, K. Yoshida, T. Morita, H. Tanaka, S.-I. Inoue, H. Yonekawa, J.-I. Hayashi, Mitochondria-related male infertility. Proceedings of the National Academy of Sciences 103, 15148–15153 (2006).

13. S. Vertika, K. K. Singh, S. Rajender, Mitochondria, spermatogenesis, and male infertility – An update. Mitochondrion 54, 26–40 (2020).

14. Z. Zhang, J. Miao, Y. Wang, Mitochondrial regulation in spermatogenesis. Reproduction, REP-21-0431 (2022).

15. J. Ramalho-Santos, S. Varum, S. Amaral, P. C. Mota, A. P. Sousa, A. Amaral, Mitochondrial functionality in reproduction: from gonads and gametes to embryos and embryonic stem cells. Human Reproduction Update 15, 553–572 (2009).

16. C. De Martino, A. Floridi, M. L. Marcante, W. Malorni, P. S. Barcellona, M. Bellocci, B. Silvestrini, Morphological, histochemical and biochemical studies on germ cell mitochondria of normal rats. Cell and Tissue Research 196, 1–22 (1979).

17. T. Tatsuta, T. Langer, AAA proteases in mitochondria: diverse functions of membrane-bound proteolytic machines. Research in Microbiology 160, 711–717 (2009).

18. P. I. Hanson, S. W. Whiteheart, AAA+ proteins: have engine, will work. Nature Reviews Molecular Cell Biology 6, 519–529 (2005).

19. T. A. Baker, R. T. Sauer, ClpXP, an ATP-powered unfolding and protein-degradation machine. Biochimica et Biophysica Acta (BBA) - Molecular Cell Research 1823, 15–28 (2012).

20. I. Levchenko, M. Yamauchi, T. A. Baker, ClpX and MuB interact with overlapping regions of Mu transposase: implications for control of the transposition pathway. Genes & development 11, 1561–1572 (1997).

21. J. Kirstein, A. Hoffmann, H. Lilie, R. Schmidt, H. Rübsamen-Waigmann, H. Brötz-Oesterhelt, A. Mogk, K. Turgay, The antibiotic ADEP reprogrammes ClpP, switching it from a regulated to an uncontrolled protease. EMBO Molecular Medicine 1, 37–49 (2009).

22. T. Wang, E. Babayev, Z. Jiang, G. Li, M. Zhang, E. Esencan, T. Horvath, E. Seli, Mitochondrial unfolded protein response gene Clpp is required to maintain ovarian follicular reserve during aging, for oocyte competence, and development of pre-implantation embryos. Aging Cell 17, e12784 (2018).

23. E. Esencan, Z. Jiang, T. Wang, M. Zhang, G. Soylemez-Imamoglu, E. Seli, Impaired Mitochondrial Stress Response due to CLPP Deletion Is Associated with Altered Mitochondrial Dynamics and Increased Apoptosis in Cumulus Cells. Reprod Sci 27, 621–630 (2019).

24. E. Esencan, M. Cozzolino, G. Imamoglu, E. Seli, Mitochondrial Stress Response Gene Clpp Is Not Required for Granulosa Cell Function. Antioxidants 10, 10.3390/antiox10010001 (2021).

25. S. Gispert, D. Parganlija, M. Klinkenberg, S. Drose, I. Wittig, M. Mittelbronn, P. Grzmil, S. Koob, A. Hamann, M. Walter, F. Buchel, T. Adler, M. Hrabe De Angelis, D. H. Busch, A. Zell, A. S. Reichert, U. Brandt, H. D. Osiewacz, M. Jendrach, G. Auburger, Loss of mitochondrial peptidase Clpp leads to infertility, hearing loss plus growth retardation via accumulation of CLPX, mtDNA and inflammatory factors. Human Molecular Genetics 22, 4871–4887 (2013).

26. Z. Pan, H. Xu, Y. Tian, D. Liu, H. Liu, R. Li, Q. Dou, B. Zuo, R. Zhai, W. Tang, W. Lu, Perrault syndrome: Clinical report and retrospective analysis. Molecular Genetics & Genomic Medicine 8, (2020).

27. F. Forli, L. Bruschini, B. Franciosi, R. Battini, G. Marinella, S. Berrettini, F. Lazzerini, A Rare Case of Perrault Syndrome with Auditory Neuropathy Spectrum Disorder: Cochlear Implantation Treatment and Literature Review. Audiology Research 11, 609–617 (2021).

28. J. C. Whitman, B. H. Paw, J. Chung, The role of ClpX in erythropoietic protoporphyria. Hematology, Transfusion and Cell Therapy 40, 182–188 (2018).

29. H. K. Gumber, J. F. McKenna, A. F. Tolmie, A. M. Jalovec, A. C. Kartick, K. Graumann, H. W. Bass, MLKS2 is an ARM domain and F-actin-associated KASH protein that functions in stomatal complex development and meiotic chromosome segregation. Nucleus 10, 144–166 (2019).

30. R. Liu, S. D. Kasowitz, D. Homolka, N. A. Leu, J. T. Shaked, G. Ruthel, D. Jain, H. Lin, S. Keeney, M. Luo, R. S. Pillai, P. J. Wang, YTHDC2 is essential for pachytene progression and prevents aberrant microtubule-driven telomere clustering in male meiosis. Cell Reports 37, 110110 (2021).

31. A. Y. Choo, S.-O. Yoon, S. G. Kim, P. P. Roux, J. Blenis, Rapamycin differentially inhibits S6Ks and 4E-BP1 to mediate cell-type-specific repression of mRNA translation. Proceedings of the National Academy of Sciences 105, 17414–17419 (2008).

32. P. M. Motta, S. A. Nottola, S. Makabe, R. Heyn, Mitochondrial morphology in human fetal and adult female germ cells. Human Reproduction 15, 129–147 (2000).

33. N. Charmpilas, N. Tavernarakis, Mitochondrial maturation drives germline stem cell differentiation in Caenorhabditis elegans. Cell Death & Differentiation 27, 601–617 (2020).

34. X. Wang, Y. Wen, J. Zhang, G. Swanson, S. Guo, C. Cao, S. A. Krawetz, Z. Zhang, S. Yuan, MFN2 interacts with nuage-associated proteins and is essential for male germ cell development by controlling mRNA fate during spermatogenesis. Development 148, dev.196295 (2021).

35. P. I. Sadate-Ngatchou, C. J. Payne, A. T. Dearth, R. E. Braun, Cre recombinase activity specific to postnatal, premeiotic male germ cells in transgenic mice. genesis 46, 738–742 (2008).

36. T. Wang, Y. Xiao, Z. Hu, J. Gu, R. Hua, Z. Hai, X. Chen, J. V. Zhang, Z. Yu, T. Wu, W. S. B. Yeung, K. Liu, C. Guo, MFN2 Deficiency Impairs Mitochondrial Functions and PPAR Pathway During Spermatogenesis and Meiosis in Mice. Frontiers in Cell and Developmental Biology 10, (2022).

37. G. Varuzhanyan, R. Rojansky, M. J. Sweredoski, R. L. J. Graham, S. Hess, M. S. Ladinsky, D. C. Chan, Mitochondrial fusion is required for spermatogonial differentiation and meiosis. Elife 8, (2019).

38. M. Li, J. Zheng, G. Li, Z. Lin, D. Li, D. Liu, H. Feng, D. Cao, E. H. Y. Ng, R. H. W. Li, C. Han, W. S. B. Yeung, L. T. Chow, H. Wang, K. Liu, The male germline-specific protein MAPS is indispensable for pachynema progression and fertility. Proc Natl Acad Sci U S A 118, (2021).

39. C. R. Clapier, B. R. Cairns, The Biology of Chromatin Remodeling Complexes. Annual Review of Biochemistry 78, 273–304 (2009).

40. H. Scherthan, S. Weich, H. Schwegler, C. Heyting, M. Härle, T. Cremer, Centromere and telomere movements during early meiotic prophase of mouse and man are associated with the onset of chromosome pairing. Journal of Cell Biology 134, 1109–1125 (1996).

41. H. Shibuya, K.-i. Ishiguro, Y. Watanabe, The TRF1-binding protein TERB1 promotes chromosome movement and telomere rigidity in meiosis. Nature Cell Biology 16, 145–156 (2014).

42. H. J. Kim, C. Liu, A. F. Dernburg, How and Why Chromosomes Interact with the Cytoskeleton during Meiosis. Genes 13, 901 (2022).

43. P. I. Sadate-Ngatchou, C. J. Payne, A. T. Dearth, R. E. Braun, Cre recombinase activity specific to postnatal, premeiotic male germ cells in transgenic mice. genesis 46, 738–742 (2008).

44. H. Gu, Y.-R. Zou, K. Rajewsky, Independent control of immunoglobulin switch recombination at individual switch regions evidenced through Cre-<em>loxP</em>-mediated gene targeting. Cell 73, 1155–1164 (1993).

45. L. Niel, D. M. Weary, Rats avoid exposure to carbon dioxide and argon. Applied Animal Behaviour Science 107, 100–109 (2007).

46. S. La Salle, F. Sun, M. A. Handel, “Isolation and Short-Term Culture of Mouse Spermatocytes for Analysis of Meiosis” (Humana Press, 2009), pp. 279–297.

47. H. Gan, L. Wen, S. Liao, X. Lin, T. Ma, J. Liu, C.-X. Song, M. Wang, C. He, C. Han, F. Tang, Dynamics of 5-hydroxymethylcytosine during mouse spermatogenesis. Nature Communications 4, (2013).

48. M. Li, J. Zheng, G. Li, Z. Lin, D. Li, D. Liu, H. Feng, D. Cao, E. H. Y. Ng, R. H. W. Li, C. Han, W. S. B. Yeung, L. T. Chow, H. Wang, K. Liu, The male germline-specific protein MAPS is indispensable for pachynema progression and fertility. Proc Natl Acad Sci U S A 118, (2021).

