## Supplementary material for "ClpP/ClpX deficiency impairs mitochondrial functions and mTORC1 signaling during spermatogenesis and meiosis": methods and materials, supplementary figures 1-4, tables 1-3

**This PDF file includes:**

Materials and Methods

Figs. S1 to S4

Tables S1 to S3

Materials and Methods

**Mouse maintenance**

All the mice used were of the C57BL/6J background and were kept in the animal facilities at Peking University Shenzhen Hospital. The animals were housed under a controlled environment with free access to water and food, and with lights switched on between 6:00 and 18:00. All experimental protocols were approved by the ethics committee of the University of Hong Kong-Shenzhen Hospital.

**Generation of the *Clpp / Clpx* conditional knock-out mouse line and genotyping**

Exon 3-exon 5 of the *Clpp* gene and exon 3 of the *Clpx* gene were targeted for deletion by insertion of two LoxP sites in the mouse genome via the CRISPR/Cas9 and homology directed repair (HDR) techniques (Fig. 1A). Two guide RNAs (gRNA), donor vector containing the LoxP sites, and Cas9 mRNA were co-injected into fertilized mouse eggs to generate the targeted cKO offspring. F_0_ founder animals were identified by PCR followed by sequencing analysis. The mice were then bred with WT mice to generate the F_1_ generation and to test for germline transmission. The gRNA sequences used are listed in supplementary table 3.

The *Clpp / Clpx* floxed mice were identified by PCR analysis with the Rapid Taq polymerase (Cat. P222-02, 2X Rapid Taq Master Mix, Vazyme, Nanjing, China). A small tissue piece was excised at the tail tip of the mice and was lysed with 200 μL of digestion buffer (50 mM KCl, 10 mM pH 9.0 Tris-HCl, 0.1 % Triton X-100 and 0.4 mg/mL Proteinase K) at 55°C overnight. Then, each sample was incubated at 98°C for 10 min to denature the Proteinase K and centrifuged at 12,000 rpm for 5 min. The supernatant containing the genomic DNA was then used in the PCR assay. The PCR primers used are listed in supplementary table 3.

The PCR mixture consisted of 1 μL of mouse tail genomic DNA, 1 μL of forward primer (10 μM), 1μL of reverse primer (10 μM), 12.5 μL of 2X Rapid Taq Master Mix, and 9.5 μL of ddH_2_O. The PCR program involved an initial denaturation step at 95°C for 3 min followed by 32 cycles comprising denaturation at 95°C for 15 s, annealing at 60°C for 15 s, and extension at 72°C for 15 s. This was followed by a final additional extension step at 72°C for 5 min. The PCR products were separated by gel electrophoresis using a 1.5% agarose gel. The WT mice showed a single band, heterozygous mice (i.e., *Clpp*^loxP/+^ *or Clpx*^loxP/+^) showed two bands, and homozygous mice (*Clpp*^lox/lox^ or *Clpx*^loxP/loxP^) demonstrated a single band with larger size (Fig. 1Bi-Bii). The homozygous ClpP/ClpX floxed mice were then crossed with the *Stra8-Cre* mice, which expressed Cre specifically in the male germ cells (*43*). Further genetic screening was conducted to select those that also expressed Cre, as shown by an additional band in gel electrophoresis (Fig. 1B), whereas the control mice did not express Cre. The ClpP/ClpX cKO mice can be identified via the Cre-LoxP system (*44*). The sequence of primers for the PCR analysis are listed in supplementary table 3.

In all experiments, the control mice were chosen from the same littermate of the cKO mice. The control mice were genotyped as *Clpp^fl/fl^*, *Clpx^fl/fl^*; homozygous ClpP/ClpX cKO mouse were genotyped as *Clpp^fl/fl^;Stra8-Cre and Clpx^fl/fl^;Stra8-Cre.*

**Tissue collection and histological analysis**

Male mice were euthanized by carbon dioxide inhalation (*45*). The testes were immediately collected and fixed in Bouin's solution (Cat. HT10132, Sigma-Aldrich, MO, USA) for 16 h before dehytodration, paraffin embedding, sectioning at 7 μm thick and mounting on glass slides. For histological examination, the tissue sections were de-paraffinized, stained with hematoxylin solution for 90 sec, washed three times with ddH_2_O, mounted with neutral balsam (Cat. G8590, Solarbio, Beijing, China) and imaged under a light microscope. For indirect immunofluorescence assays, the testes were fixed in 4% paraformaldehyde for 24 h before processing as described above.

**Chromosome spread assays**

The tunica albuginea of the testes was removed and the testicular tissue was transferred into a 1.5 mL RNase/DNase Free tube (Cat. AXYMCT150CS, Axygen, Corning, NY, USA) containing 1 mL PBS (Cat. 10010023, Gibco, Thermo Fisher Scientific, MA, USA). A pair of forceps (Cat. HEC7.1, Tweezers Round, Carl Roth, Karlsruhe, Germany) were used to crush the seminiferous tubules for around 5 min, or until the large tissue pieces had been dissociated. The resulting cell suspension was filtered into a new 1.5 mL microtube though a 40 μm cell strainer (Cat. 352340, Falcon, Corning, NY, USA). The cells were centrifuged at 3,500 rpm for 3 min, after which the supernatant was discarded. The pellet was gently resuspended in 1 mL PBS and the cells were washed by centrifugation 1-2 times until the supernatant was clear. After the final wash, the cells were resuspended in 1 mL of hypotonic solution (30 mM Tris HCl, 17 mM trisodium citrate, 5 mM EDTA, 50 mM sucrose, pH 8.8) and incubated at room temperature for 30-40 min. The cells were centrifuged at 3,500 rpm for 3 min and the cell pellet was resuspended and incubated in 100 μL sucrose (100 mM) for 5 min at room temperature before fixation by addition of same volume of fixative buffer (0.33 M paraformaldehyde, 100 μL of 10% Triton X-100, 10 mL PBS and 30 μL of 1N NaOH). Two circles were drawn on glass slides with a PAP pen (Cat. Z377821, Sigma-Aldrich) to mark the location of the specimen, after which 20 μL of the fixed cell suspension was applied to each circle, and the cells were allowed to settle and attach for 3 h at room temperature. Once the cell suspension was completely dry, the slides were stored at -80°C for up to 3 months prior to the immuno-staining.

**Immunofluorescence assay**

Fixed cells or tissue sections were permeabilized in phosphate buffered saline (PBS) containing 0.1% Triton X-100 (Cat. T8787, Sigma-Aldrich; PBST) for ~15 min. The cells/tissues were washed twice with PBST, after which they were incubated with blocking PBS buffer containing 5% BSA (Cat. A1933, Sigma-Aldrich, MO, USA) for 1 h at room temperature. The samples were then incubated with the primary antibody in blocking buffer overnight at 4°C, before washing for 10 min with three changes of PBS and incubation with appropriate Alexa Fluor-tagged secondary antibody (1:500 dilution in blocking buffer) for 1 h at room temperature. The samples were then washed thoroughly with PBST and briefly with distilled H_2_O, after which they were mounted with the ProLong Diamond Antifade mountant containing DAPI (Cat. P36962, Invitrogen, MA, USA). The samples were incubated at room temperature overnight to cure the mountant, and then stored at 4°C prior to examination under a confocal microscope. The antibodies and their dilution used are listed in Supplementary Table 1 and 2.

Mitochondrial membrane potential was evaluated using the JC-1 probe (Cat. T3168, Invitrogen, MA, USA). In brief, spermatocytes were cultured in DMEM/F12 medium containing 2 mM JC-1 for 30 min at 37°C and washed for 3 x 3 min with PBS. The samples were immediately imaged by confocal microscopy. JC-1 dye exhibits a potential-dependent accumulation in mitochondria as indicated by an emission shift of fluorescence from green (~529 nm) to red (~590 nm). Thus, mitochondrial depolarization is indicated by a decrease in the red/green fluorescence intensity ratio.

Reactive oxygen species (ROS) in spermatocytes were determined by carboxy-H2DCFDA (Cat. C400, Invitrogen, MA, USA), which is a fluorescent oxidative stress indicator (Takeda et al*.*, 2003). The spermatocytes were pre-treated with DMEM/F12 medium containing 10 mM H_2_O_2_ for 5 min. They were then washed and incubated with 10 µM carboxy-H2DCFDA in DMEM/F12 for 30 min at 37°C, after which they were washed 3 times each for 3 min with PBS and then immediately imaged under a confocal microscope.

**Confocal imaging**

Images of the fluorescence labelled tissue sections or cells were acquired using a ZEISS LSM 900 with Airyscan 2 laser scanning confocal microscope with Hybrid Detectors (HyD). Images were captured with either a Leica HC PL APO CS2 63x/1.4 NA oil immersion objective lens or a Leica HC PL APO 20x/0.7 NA CS2 dry objective lens. Alex Fluor 488, Alex Fluor 546/594, Alex Fluor 647 and DAPI fluorescence were captured with an argon laser operating at 488 nm, a HeNe laser at 561 nm, a HeNe laser at 633 nm, or a diode-pumped solid-state laser at 405 nm, respectively, using 488 nm excitation/519 nm detection, 552 nm excitation/575 nm detection, 633 nm excitation/670 nm detection, and 405 nm excitation/461 nm detection, respectively.

**Purification of male germ cells**

Male mice germ cells were purified following a well-establish protocol (*46,47,48*). In brief, mice were sacrificed, and the testes were decapsulated. The seminiferous tubules were isolated from other testicular tissues by incubation with 0.2% W/V collagenase type I (Cat. 17100017, Gibco, MA, USA) and 1 mg/mL DNase I (Cat. A3778, Applichem, IOWA, USA) in a water bath at 37°C for 10 min, after which the samples were centrifuged at 1200 rpm for 2 min. The pellet containing the seminiferous tubules was resuspended in 5 ml of 0.25% trypsin (Cat.15050057, Gibco, MA, USA) containing 1 mg/mL DNase I at 37°C for 5 min with gentle shaking. The sample was centrifuged at 1500 rpm for 5 min, after which the pelleted cells were resuspended in 20 mL of high glucose DMEM (Cat. 12100046, Gibco, MA, USA) containing 0.5% BSA and filtered through a 40 μm nylon cell strainer (Cat. 352340, Falcon®, BD, NJ, USA). The recovered cells in the filtrate were resuspended in 20 mL DMEM containing 0.5% BSA and loaded into a cell separation apparatus (BOMEX Corporate) containing a 2%–4% BSA gradient in 600 mL of DMEM. After sedimentation for 3 h, the cells were collected into tubes from the bottom of the separation apparatus at a rate of 10 mL/min. The cell type and purity in each fraction were assessed according to their diameter and morphological characteristics under a light microscope.

**Protein extraction and western blotting analysis**

Purified spermatocytes from WT and ClpP/ClpX cKO male C57BL/6 mice testes were suspended in lysis buffer [50 mM HEPES-KOH (pH 7.5), 100 mM KCl, 2 mM EDTA, 10% glycerol, 0.1% NP-40, 10 mM NaF, 0.25 mM Na_3_VO_4_, and 50 mM β-glycerophosphate] supplemented with complete protease inhibitor (Cat. 04693116001, Roche, Basel, Switzerland). The samples were homogenized and centrifuged at 20,000 g for 20 min at 4°C, after which the supernatant was retained for western blotting analysis. The proteins in each sample were separated using 4%-12% Bis-Tris gels (Cat. M00652, SurePAGE™, GenScript, Nanjing, China) and a mini protein electrophoresis system (Cat. 1658034, BIO-RAD, CA, USA) following the manufacturer’s instructions. The protein bands were transferred to polyvinylidene fluoride (PVDF) membranes (Cat. IPVH00010, Immobilon, Millipore, MA, USA) via a Mini Trans-Blot Electrophoretic Transfer Cell (Cat. 1703930, BIO-RAD, CA, USA). The immunoreactive bands were detected and analyzed with a BIO-RAD ChemiDoc MP imaging System (Cat. 12003154, BIO-RAD, CA, USA) in conjunction with the Image Lab Software (BIO-RAD, CA, USA). The relative protein levels in each sample were normalized to β-tubulin or β-actin to standardize the loading. The primary and secondary antibodies for immunoblotting are shown in supplementary table 1 and 2.

**Quantitative reverse-transcription polymerase chain reaction (qRT-PCR)**

Total RNA was extracted from purified pachytene spermatocytes using BSA gradient sedimentation and was reverse transcribed using the High-capacity cDNA Reverse Transcription kit (Cat. 4368813, Thermo Fisher Scientific, CA, USA) according to the manufacturer’s instructions. Five genes (*Sdhb*, *Uqcrc2*, *Atp5a1*, *Ndufv1* and *Cox1*) and the house-keeping gene *Gapdh* were quantified using the real time-qPCR with SYBR® Green master mix (Cat. 1725124, iTaq™ Universal SYBR® Green Supermix, BIO-RAD, CA, USA), and an Applied Biosystems 7500 Real-time PCR system (Cat. 4351107, Applied Biosystems™, Thermo Fisher Scientific, MA, USA). The following program was used: Activation at 95°C for 3 min (1 cycle) and then 40 cycles of denaturation at 95°C for 15 sec and annealing/extension at 60°C for 30 sec. The data were normalized to GAPDH and the relative levels of mRNA were quantified using the 2^-ΔΔCt^ method. The primers used are listed in supplementary table 3.

**RNA sequencing and data analysis**

RNA sequencing was performed in an Illumina HiSeq PE150 (Novogene Corp. Inc., Beijing, China). Total RNA was purified from pachytene spermatocytes using the Trizol reagent (Cat. 15596026, TRIzol™, Thermo Fisher Scientific, CA, USA) according to the manufacturer’s instructions. The RNA quality and quantity were evaluated using a Bioanalyzer (Agilent Co.) and a Qubit (Life Technologies), respectively, and normalization was performed using the Novomagic software (Novogene Corp. Inc.) to generate the number of reads in fragments per kilobase per million mapped fragments (FPKM). Genes with an average FPKM > 1 in at least one condition were used in subsequent scatterplot analyses. Differential expression analysis was performed using the DESeq2, and genes with an adjusted *P*-value < 0.05 were considered as being differentially expressed. Expression pattern clusters were generated by unsupervised hierarchical clustering analysis and K-means clustering algorithm using the R. DAVID, and the Ingenuity Pathways Analysis (IPA) were used to reveal the Gene Ontology (GO) and Kyoto Encyclopedia of Genes and Genomes (KEGG) analysis.

**Methylated RNA immunoprecipitation (MeRIP) sequencing and data analysis**

The mRNA of m6A was sequenced by the MeRIP-seq at Novogene (Beijing, China). Briefly, a total of 300 µg RNA was extracted from pachytene spermatocytes. The integrity and concentration of the extracted RNA was determined in an Agilent 2100 bioanalyzer (Agilent) and a simpliNano spectrophotometer (GE Healthcare), respectively. Fragmented mRNA (~100 nt) was incubated for 2 hr at 4℃ with anti-m6A polyclonal antibody (Cat. 202-003, Synaptic Systems, Göttingen, Germany) in the immunoprecipitation experiment. The immunoprecipitated mRNAs or Input was used for library construction with the NEB Next ultra-RNA library prepare kit for Illumina (New England Biolabs). The libraries were sequenced on an Illumina Novaseq or Hiseq platform with a paired-end read length of 150 bp according to the standard protocols. The sequencing was carried out with 30 independent biological replicates. Raw data of fastq format were firstly processed using the fastp (version 0.19.11). Reference genome and gene model annotation files were downloaded from the NCBI genome website (https://www.ncbi.nlm.nih.gov). After mapping reads to the reference genome, the exomePeak R package (version 2.16.0) was used for the m6A peak identification in each anti-m6A immunoprecipitation group with the corresponding input samples serving as a control, and a q-value threshold of enrichment of 0.05 was used for all data sets. Using the differential peak calling, genes associated with differential peaks were identified followed by GO and KEGG analyses.

**Drug preparation and *in vivo* pharmacological treatment**

Rapamycin (Cat. HY-10219-25mg, MCE, New Jersey, United States) was firstly dissolved in 1 mL DMSO (Cat. D2650, Sigma, St. Louis, United States) with slight vortexing to prepare a stock concentration of 25 mg/mL. The stock rapamycin solution was stored at -80°C up to 6 months. Before injection, the stock solution was diluted with PBS to a final injection concentration of 3.125 mg/mL. The ClpP cKO mice and ClpX cKO mice were injected intraperitoneally every other day with 8 mg/kg rapamycin, from PD 10 to PD 35. The control mice from same litter were injected with the same volume of PBS. After rapamycin treatment, the mice were sacrificed and their testicular tissue were collected for molecular and histological evaluation.

**Statistical analysis**

The data were analyzed with the Minitab version 18 (Minitab Inc. State College, PA, USA) using the Student’s *t*-test or one-way ANOVA followed by Tukey’s test. *P*-values less than 0.05 were considered to be statistically significant. Graphs were generated using Microsoft Excel and figures were prepared with the CorelDraw version X8 (Corel Corp., Ottawa, ON, Canada).

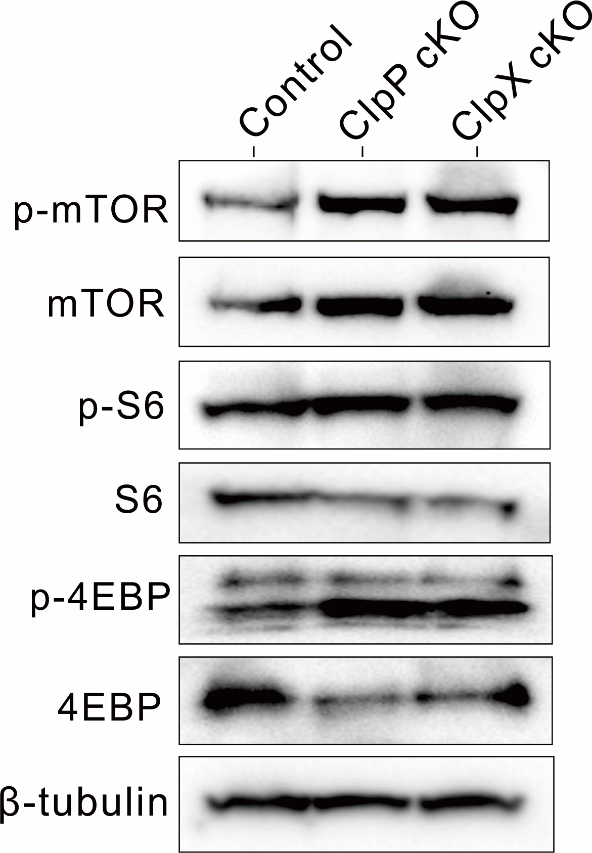

Fig. S1. The mTORC1 signaling was activated in ClpP/ClpX cKO spermatocytes. The spermatocytes from control mice and ClpP/ClpX cKO mice were isolated before the proteins were extracted. The protein expression level of pan-mTOR, phospho-mTOR, and two substrates of mTORC1 signals, including S6 and 4EBP, as well as their phosphorylated proteins were determined via western blot analysis. β-tubulin acts as a marker for controlling the protein loading for each lane.

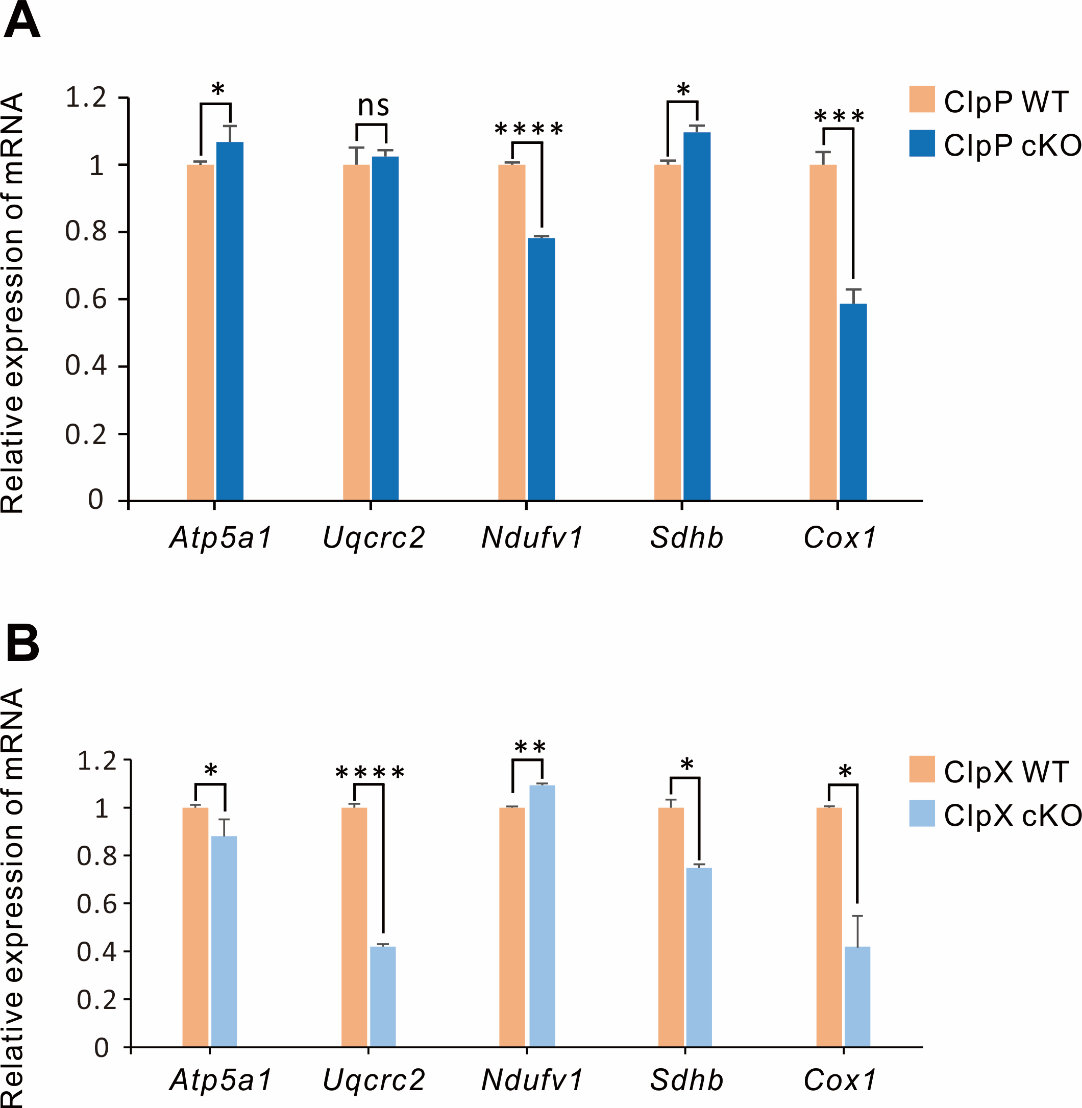

Fig. S2. The mRNA level of respiratory chain genes in ClpP/ClpX cKO spermatocytes. The spermatocytes from (A) *Clpp^fl/fl^* mice and *Clpp^fl/fl^;S8-Cre* mice or (B) *Clpx^fl/fl^* mice and *Clpx^fl/fl^;S8-Cre* mice were isolated before the RNA were extracted. The mRNA expression levels were measured by RT-qPCR assay. *: p<0.05, **: p<0.01, ***: p<0.001.

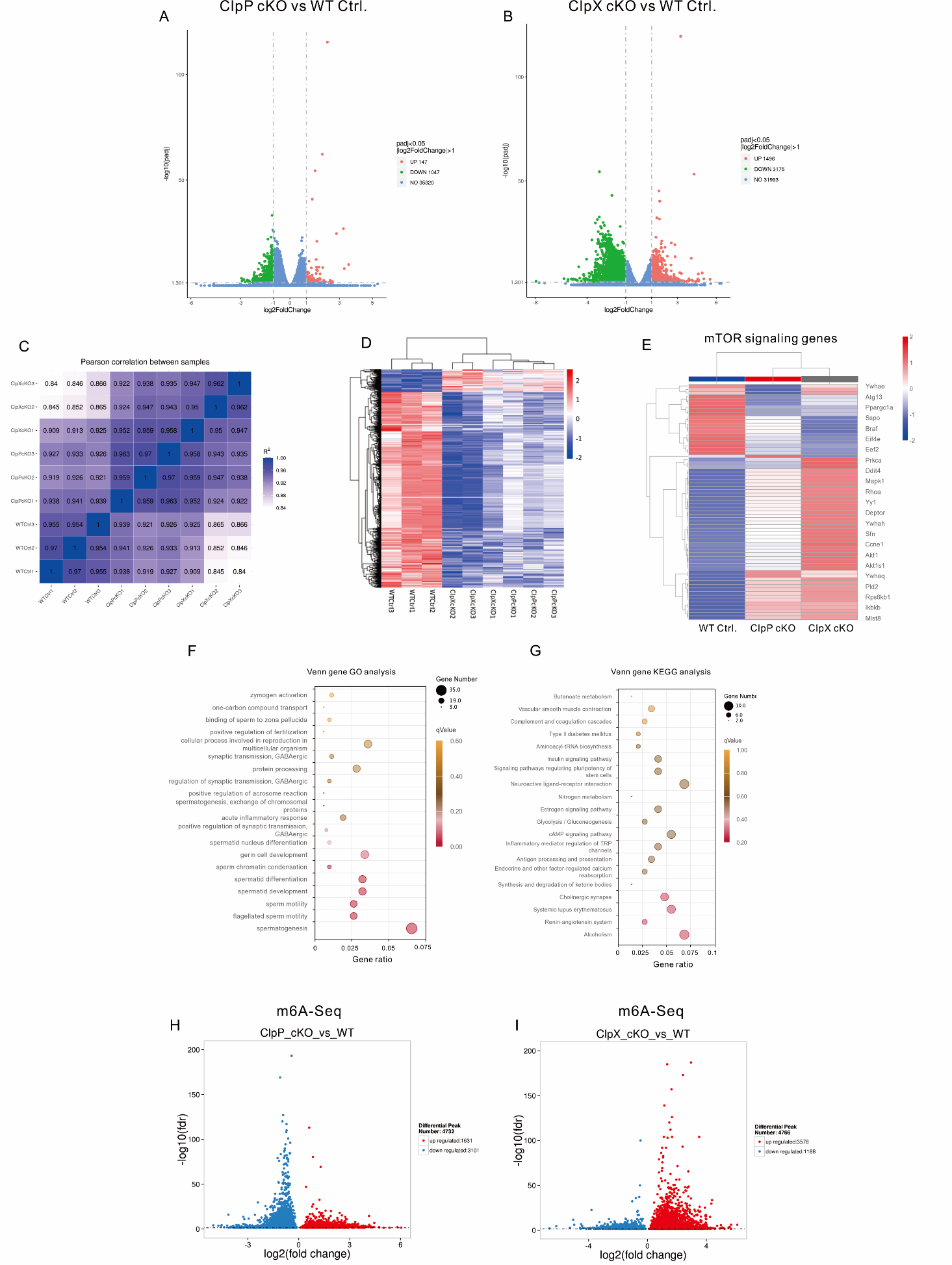

Fig. S3. The RNA sequencing and m6A sequencing analysis of spermatocytes in WT control, ClpP cKO and ClpX cKO groups. (A-B) Volcano plots of RNA-seq from ClpP cKO and ClpX cKO spermatocytes comparing with controls. Green plots and red plots represent down-regulated genes and up-regulated genes, respectively. (C) Pearson correlation analysis among sequenced samples. (D) Heatmap and cluster analysis of 1005 common genes obtained from Venn analysis. (E) Heatmap and cluster analysis of mTOR signaling genes expression. (F-G) Dot plots of GO and KEGG analysis for 1005 common genes. (H-I) Volcano plots of m6A-seq data from ClpP cKO and ClpX cKO spermatocytes comparing with controls, blue plots and red plots represent down-regulated genes and up-regulated genes, respectively.

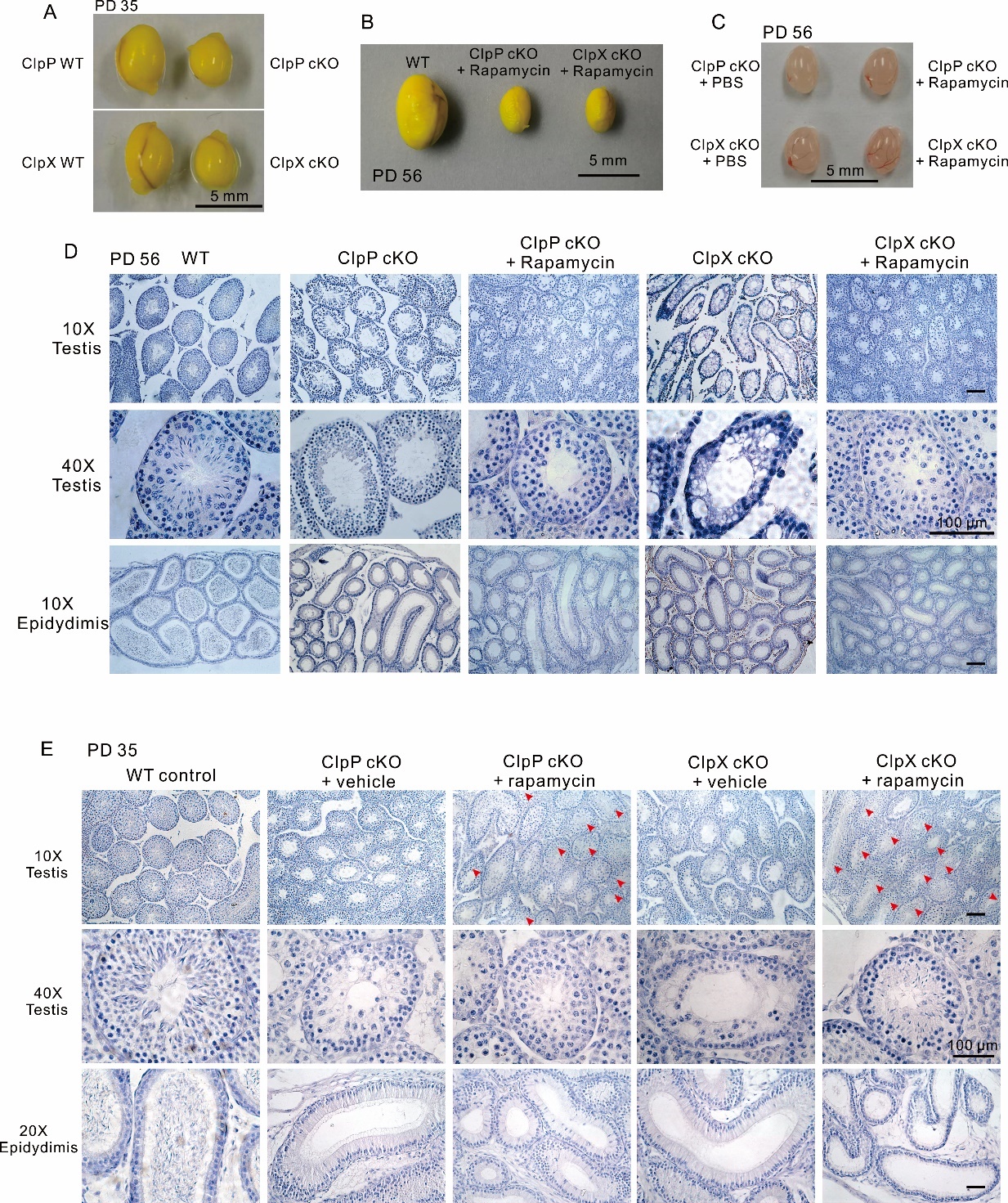

Fig. S4. Morphology analysis showed rapamycin treatment helps ClpP/ClpX cKO germ cell in meiotic process. (A) The morphology of ClpP/ClpX cKO testes and related controls at PD 35. (B) The morphology of testes from rapamycin injected ClpP/ClpX cKO mice (i.p. injection from PD 14- PD 56, every other day) and WT testis at PD 56. (C) The morphology of testes from PBS or rapamycin injected ClpP/ClpX cKO mice (i.p. injection from PD 14- PD 56, every other day) at PD 56. Scale bars are 5 mm. (D) Histological analysis of testes and epididymis from WT control, ClpP/ClpX cKO and rapamycin injected ClpP/ClpX cKO mice at PD 56. (E) Histological analysis of testes and epididymis from WT control, PBS injected ClpP/ClpX cKO and rapamycin injected ClpP/ClpX cKO mice at PD 35. Red arrow heads indicate elongated spermatozoa in seminiferous tubules of rapamycin treated ClpP/ClpX cKO mice. Scale bars are 100 μm.

Table S1.

Primary Antibody List

| Primary antibodies | Host | Dilution rate | Catalog No. | Manufacturer | Country |
| --- | --- | --- | --- | --- | --- |
| Anti-gamma H2A.X (phosphor Ser139) | Rabbit | 1:300 | ab81299 | Abcam | Cambridge, UK |
| Anti-Sycp3 | mouse | 1:300 | ab181746 | Abcam | Cambridge, UK |
| Anti-cleaved-PARP | Rabbit | 1:200 | #94885 | Cell Signaling Technology | MA, USA |
| Anti-H1t | Rat | 1:100 | --------- | Custom made by Dai·An | Wuhan, China |
| Anti-cleaved-caspase 3 | Rabbit | 1:200 | #9664 | Cell Signaling Technology | MA, USA |
| Anti-TRF1 | Rat | 1:200 | ab192629 | Abcam | Cambridge, UK |
| Anti-MLH1 | Mouse | 1:200 | 554073 | BD | NJ, USA |
| Anti-alpha Tubulin | Rabbit | 1:300 | ab52866 | Abcam | Cambridge, UK |
| Anti-beta Tubulin | Rabbit | 1:300 | #2146S | Cell Signaling Technology | MA, USA |
| Anti-phospho S6 | Rabbit | 1:1000 | #4856 | Cell Signaling Technology | MA, USA |
| Anti-S6 | Rabbit | 1:1000 | #2217 | Cell Signaling Technology | MA, USA |
| Anti-phospho S6K | Rabbit | 1:300 | #9234 | Cell Signaling Technology | MA, USA |
| Anti-S6K | Rabbit | 1:1000 | #2708 | Cell Signaling Technology | MA, USA |
| Anti-phospho 4EBP1 | Rabbit | 1:1000 | #2855 | Cell Signaling Technology | MA, USA |
| Anti-4EBP1 | Rabbit | 1:1000 | #9452 | Cell Signaling Technology | MA, USA |
| Anti-phospho mTOR (Ser2481) | Rabbit | 1:1000 | #2974 | Cell Signaling Technology | MA, USA |
| Anti-mTOR | Rabbit | 1:1000 | #2983 | Cell Signaling Technology | MA, USA |
| Anti-β-tubulin | Rabbit | 1:2000 | #2146 | Cell Signaling Technology | MA, USA |
| Anti-ClpP | Rabbit | 1:1000 | #14181 | Cell Signaling Technology | MA, USA |
| Anti-ClpX | Rabbit | 1:1000 | ab168338 | Abcam | Cambridge, UK |

Table S2.

Secondary Antibody List

| Secondary antibodies | Host | Dilution rate | Cat | Manufacturer | Country |
| --- | --- | --- | --- | --- | --- |
| Alexa Fluor Plus 488-tagged goat anti-mouse IgG (H+L) highly cross-adsorbed antibody | Goat | 1:500 | A32723 | Invitrogen | MA, USA |
| Alexa Fluor Plus 488-tagged goat anti-rabbit IgG (H+L) highly cross-adsorbed antibody | Goat | 1:500 | A32731 | Invitrogen | MA, USA |
| Alexa Fluor Plus 594-tagged goat anti-mouse IgG (H+L) highly cross-adsorbed antibody | Goat | 1:500 | A32742 | Invitrogen | MA, USA |
| Alexa Fluor Plus 594-tagged goat anti-rabbit IgG (H+L) highly cross-adsorbed antibody | Goat | 1:500 | A32740 | Invitrogen | MA, USA |
| Alexa Fluor Plus 488-tagged goat anti-rat IgG (H+L) highly cross-adsorbed antibody | Goat | 1:500 | A48262 | Invitrogen | MA, USA |
| HRP conjugated goat anti-mouse IgG (H+L) cross-adsorbed antibody | Goat | 1:5000 | G-21040 | Invitrogen | MA, USA |
| HRP conjugated goat anti-rabbit IgG (H+L) cross-adsorbed antibody | Goat | 1:5000 | G-21234 | Invitrogen | MA, USA |

Table S3.

| Name | Sequence (PAM region underlined) |
| --- | --- |
| *Clpp* gRNA1 | 5’-AAATGTCCTGTCAGCTACATGGG-3’ |
| *Clpp* gRNA2 | 5’-CATAGAACCTATGTCGGGCCAGG-3’ |
| *Clpx* gRNA1 | 5’-GTTACTTATAGGAATAATCCAGG-3’ |
| *Clpx* gRNA2 | 5’-CCGTGGAGCTCAAGGGCAGTTGG-3’ |
| *Clpp* forward primer | 5’- TCTTACCATAGTCTGCTGTTGTCA-3’ |
| *Clpp* reverse primer | 5’- CTACACCTGGTCTGGTTTAGGAAC-3’ |
| *Clpx* forward primer | 5’- GTAAGTCTCCTGATCCAACCT-3’ |
| *Clpx* reverse primer | 5’- TGTCCAGCTAAGGATTCTCATT-3’ |
| *Stra8* forward primer | 5’-GTGCAAGCTGAACAACAGGA-3’ |
| *Stra8* reverse primer | 5’- AGGGACACAGCATTGGAGTC-3’ |
| *Sdhb* forward primer | 5’-AATTTGCCATTTACCGATGGGA-3’ |
| *Sdhb* reverse primer | 5’-AGCATCCAACACCATAGGTCC-3’ |
| *Uqcrc2* forward primer | 5’-AAAGTTGCCCCGAAGGTTAAA-3’ |
| *Uqcrc2* reverse primer | 5’-AAAGTTGCCCCGAAGGTTAAA-3’ |
| *Atp5a1* forward primer | 5’-TCTCCATGCCTCTAACACTCG-3’ |
| *Atp5a1* reverse primer | 5’-CCAGGTCAACAGACGTGTCAG-3’ |
| *Ndufv1* forward primer | 5’-TTTCTCGGCGGGTTGGTTC-3’ |
| *Ndufv 1* reverse primer | 5’-GGTTGGTAAAGATCCGGTCTTC-3’ |
| *Cox1* forward primer | 5’-GTGCTGGGGCAGTGCTGGAG-3’ |
| *Cox1* reverse primer | 5’-TGGGGCCTGAGTAGCCCGTG-3’ |
| *Gapdh* forward primer | 5’-GGCAAATTCAACGGCACAGT-3’ |
| *Gapdh* reverse primer | 5’-GGCCTCACCCCATTTGATGT-3’ |

The sequence of gRNAs and primers
